# Combining CHARMM36m with OPC water improves accuracy

**DOI:** 10.64898/2026.09.25.754057

**Authors:** Nicolai Kozlowski, Gabor Nagy, Crystal F. Ottoway, Lars V. Bock, Sara M. Vaiana, Helmut Grubmüller

## Abstract

Atomistic simulations of intrinsically disordered proteins (IDPs) are notoriously sensitive to force field inaccuracies, either regarding the protein or the water model, yielding inaccurate observables such as compactness, secondary structure propensities, or kinetics.

The currently most widely used IDP force fields are Amber99sb-disp (A99disp) and Charmm36m (C36m). A99disp includes a new water model and thus optimized both, the protein and the water interactions. In contrast, C36m used the Tip3p water model and optimized only protein interactions. In many cases, C36m+Tip3p underestimates radii of gyration compared to FRET or SAXS experiments. Such overly compact structural ensembles are believed to arise from an imbalance between protein-protein, protein-water, and water-water interactions, which might be due to Tip3p inaccuracies.

Here, we aim at re-balancing these interactions by combining C36m with the Optimal Point Charge (OPC) water model. Recently, this C36m+OPC combination showed improved accuracy for the disordered domain of the measles virus nucleoprotein. Here we present a systematic assessment, comparing C36m+OPC, C36m+Tip3p, C36m+Tip4p, C22*, A03ws, A99sb-ws, and A99disp for five IDPs, as well as a subset of those for five globular proteins, a set of disordered AGQ-repeat peptides, and the fast folding miniprotein CLN025. We compared extensive MD simulations (*>* 8.5 ms) with SAXS, NMR, circular dichroism, photo-induced electron transfer (PET), T-jump infrared spectroscopy, and X-ray crystallography measurements.

We found that combining C36m with OPC improved accuracy over C36m+Tip3p for IDP ensembles without compromising its accuracy for globular proteins. While also the kinetics of the AGQ-peptides were more accurate for C36m+OPC, those of CLN025 folding were less accurate. Overall, C36m+OPC showed similar accuracy as A99sb-ws and A99disp, the latter is currently considered among the most accurate protein force fields.

## 1 Introduction

Accurate molecular dynamics (MD) simulations of intrinsically disordered proteins (IDPs) have proven notoriously challenging, because of their high sensitivity to force field inaccuracies.^1^ Currently, the most widely-used IDP force fields are Amber99sb-disp^2^ (A99disp), which was developed together with a new water model based on Tip4p-D,^3^ and Charmm36m^4^ (C36m), which was developed using the (Charmm-modified) Tip3p^5,6^ water model. It has been recently observed that, for certain larger IDPs, simulations using C36m+Tip3p predict markedly smaller radii of gyration (*R*_g_) than those estimated from fluorescence resonance energy transfer (FRET) experiments.^4^ This overestimation of the compactness of IDPs is a common inaccuracy of many force fields ^2–4,7–9^ and is assumed to be largely due to an inaccurate balance between protein-protein, protein-solvent, and solvent-solvent interactions.^4^ This imbalance is likely caused by older water models, such as Tip3p, which tend to underestimate the strength of protein-solvent interactions. ^2–4,8–14^

This issue was thoroughly addressed during the development of A99disp^2^ by simultaneously optimizing protein and water parameters. For C36m+Tip3p, however, the developers mainly focused on optimizing protein parameters, and only improved the Tip3p water parameters by ad-hoc scaling of the protein-water van der Waals interactions of water hydrogen atoms.^4^ Here, we aim to improve C36m by changing the water model to reduce this interaction imbalance.

Across different force fields, various adaptations were proposed to rebalance the strength of protein-solvent interaction and were tested with mixed success. These adaptations include increasing the van der Waals well depth of water oxygen atoms either for all interactions ^2,3^ or restricted to protein-water interactions,^8^ increasing the van der Waals well depth of water hydrogen atoms specifically for protein-water interactions,^4^ exchanging the water model, ^2,3,9,15^ and modifying protein parameters.^2,15^

Exchanging water models showed promising results in earlier studies, ^3,9,15^ but can introduce inaccuracies due to parameter inconsistencies.^16^ Specifically, strengthening protein-water interaction can increase the accuracy for IDPs but destabilize or perturb globular proteins,^2–4,8,15^ as was shown for the combinations of Amber99sb-ildn^17^ (A99sb-ildn) and Amber12^18^ with Tip4p-D water (replacing Tip3p).^3^ Hence, thorough validation of any new combination, using a large set of proteins and experiments, is required to ensure its accuracy.

Recently, Gong et al.^15^ aimed to increase the accuracy of C36m for IDPs by combining it with the A99disp water model (among other changes). They showed improved accuracy for the *R*_g_ and secondary structure of one IDP and tested thermodynamic stability of four globular proteins in 4 µs simulations each, but their validation lacks more direct comparisons of old and new force field combinations with experiments to thoroughly quantify their accuracies.

Here, we propose to combine C36m with a different water model, the optimal point charge water (OPC),^19^ and validate this combination. We chose the OPC water model because it accurately reproduces multipole moments of water molecules,^19^ bulk water properties,^19^ solvation free energies,^19^ and other observables relevant to simulations of biomolecules.^9,20,21^ Specifically, OPC showed an increased accuracy for IDPs in combination with A99sb-ildn.^9^

Previously, combining C36m+OPC to simulate the disordered region of the measles virus nucleoprotein (MeV-N-ail) resulted in good agreement with measurements from small angle X-ray scattering (SAXS), nuclear magnetic resonance (NMR), and circular dichroism spectroscopy (CD) experiments.^22^ Building on these results, we here assess whether C36m+OPC generally predicts IDP ensembles more accurately than C36m+Tip3p without compromising accuracy for globular proteins or for kinetics.

To this end, we benchmark C36m+OPC for five IDPs with diverse sizes, secondary structure propensities, and transient tertiary structure,^22,23^ as well as for five globular proteins covering the size range of typical small proteins, different secondary structure compositions, and different fold types. To asses force field accuracy, we compare the obtained results to SAXS, NMR, and CD experiments. To evaluate how C36m+OPC compares to other force field combinations, we assess not only the accuracy of C36m+OPC and C36m+Tip3p, but also of C36m+Tip4p,^24^ C22*,^16,25,26^ A03ws,^8,27,28^ A99sb-ws,^8,17,27,29^ A99sb-ildn^17,27,29^ and specifically the well established A99disp,^2^ which is widely considered one of the most accurate force fields for disordered and globular proteins.^2,15,30^

In addition to the thermodynamics properties probed by these ensemble-based measurements, we also assess the accuracy of protein kinetics. To this end, Cys-Trp (C-W) contact formation times observed in the simulations of disordered AGQ-repeat peptides are compared with results from photo-induced electron transfer (PET) experiments. Additionally, predicted folding kinetics of CLN025^31^ are compared with temperature-jump infrared spectroscopy (T-jump IR) experiments.^32^

Overall, C36m+OPC showed an improved accuracy over C36m+Tip3p for IDP ensembles without compromising its accuracy for globular proteins. The accuracy of kinetics improved for the AGQ-peptides but worsened for CLN025 folding. In our assessment, the most accurate force fields for IDPs were found to be C36m+OPC, A99disp, and A99sb-ws.

## 2 Methods

### 2.1 Selection of Test Proteins

To test the accuracy of seven force fields, including the new combination C36m+OPC, for IDPs, we selected a set of five IDPs which cover different sizes, transient secondary structures, and transient non-local interactions. The set contains (1) MeV-Ntail, the disordered domain of the measles virus nucleoprotein (127 residues) which shows transient *α* and *β* secondary structure and transient tertiary structure,^22^ (2) p53-TAD, the N-terminal transactivation domain of p53 (73 residues, PDB: 2L14^33^) which shows transient helical secondary and transient tertiary structure,^23^ (3) CBP-NCBD, the nuclear coactivator binding domain of the CREB binding protein (59 residues, PDB: 2L14,^33^ 1KBH,^34^ 1ZOQ^35^) which shows both stable and transient *α*-helices,^36^ (4) p53-pTAD, a 49 residue long part of p53-TAD (residues 13–61, PDB: 2L14^33^), and (5) RSP-8, an RS-repeat peptide with sequence GAMGPSYG-(RS)_8_ with very small secondary structure propensities.^36^

To test the accuracy of C36m+OPC for globular proteins, we selected a set of five globular proteins, which cover the size range of typical small proteins, different secondary structure compositions and tertiary structures, and have no prosthetic groups. This set contains (1) TIM, a homodimer of triose phosphate isomerase (*α/β* barrel) with 247 residues (PDB: 1TIM^37^), (2) lysozyme (HEWL, *α/β*) from hen egg white with 129 residues containing four disulfide bridges (PDB: 193l ^38^), (3) ribonuclease A (RNaseA, *α/β*) with 124 residues (PDB: 3RN3^39^), (4) the PDZ-domain of neuronal nitric oxide synthase (nNOS, *α/β*) with 112 residues (PDB: 1QAU ^40^), and (5) the XPC-binding domain of protein hHR23B (XPCBD, all-*α*) with 72 residues (PDB: 1PVE^41^).

To test the accuracy of kinetics predicted from the simulations, for which available experimental data is sparse, we included a set of disordered AGQ-repeat peptides, namely S-(AGQ)_5_-C-(AGQ)*_n_*-W and C-(AGQ)_5_-S-(AGQ)*_n_*-W for *n ∈ {*1, 2, 3*}*. For these peptides, we performed photoinduced electron transfer (PET) relaxation measurements that probe the C-W contact-formation time. Additionally, we included the 10-residue synthetic protein CLN025 (PDB: 5AWL^31^), for which temperature-jump infrared spectroscopy (T-jump IR) measurements of its folding kinetics are available. ^32^

### 2.2 Molecular Dynamics Simulations

We performed MD simulations for the disordered proteins with the following force fields: Charmm36m^4^ with OPC^19^ water model (C36m+OPC), the canonical Charmm36m with Charmm-modified Tip3p^5,6^ water model (C36m+Tip3p), Charmm36m with Tip4p/2005^24^ water model (C36m+Tip4p, only MeV-Ntail and RSP-8), Charmm22*^16,25,26^ with Charmm-modified Tip3p water model (C22*), Amber03-ws^8^ with Tip4p/2005 water model (A03ws), Amber99sb-disp^2^ with its specific water model (A99disp), and Amber99sb-ws^27^ with Tip4p/2005 water model (A99sb-ws).

For the AGQ-repeat peptides, we performed simulations with the C36m+OPC, C36m+Tip3p, A99disp, and A99sb-ws force fields. Globular proteins and CLN025 were simulated using C36m+OPC, C36m+Tip3p, A99disp, and Amber99sb-ildn^17^ with Tip4p-Ew^42^ water model (A99sb-ildn). Hence, except C36m+OPC and C36m+Tip4p, all force fields were used in combination with the water model recommended by the respective developers.^2,4,8,16,17,27^

We describe the general simulation procedure here, further details and MD parameters are provided in Supp. Tables S1–S3. We used Gromacs^43^ for all simulations. The starting structure for simulations of the globular proteins, CLN025, and part of the simulations of p53-TAD, CBP-NCBD, and p53-pTAD were taken from the PDB^44^ entries mentioned above. All other simulations were started from extended structures. For some of the IDP simulations and all AGQ-peptide simulations, the extended structures were partially collapsed before starting the MD simulations, using simulations in vacuum or implicit solvent (Supp. Methods). Solvent and ions (Na^+^ and Cl*^−^*) were added to establish a physiological salt concentration and to neutralize the overall system charge. Energy minimization was performed, followed by an NVT equilibration with position restraints applied to protein heavy atoms. Subsequently, for IDPs and AGQ-peptides, an NPT equilibration was performed first with position restraints (NPT1) and then without (NPT2). For CLN025 and globular proteins, NPT equilibration was only performed without position restraints. All bond lengths were constrained and hydrogen atoms were treated as virtual sites.^45^ Van der Waals forces were calculated using a cutoff, and Coulomb forces were calculated using the particle mesh Ewald method (PME).^46^

Errors of quantities calculated from MD trajectories were estimated using trajectory-wise bootstrapping. If only a single trajectory was available for a combination of protein and force field, it was divided into three parts, with subsequent bootstrapping between those parts.

### 2.3 Comparison to Experiments

#### 2.3.1 Small-Angle X-ray Scattering Intensity Curves

To validate the force fields, we compared measured SAXS intensity curves, i.e., the azimuthally averaged intensity of the scattered signal *I*(*q*) as a function of the scattering vector *q*, to those calculated from our MD trajectories. SAXS curves were calculated from the MD trajectories using CRYSOL,^47^ assuming a hydration shell thickness of 3 Å (considering only the first hydration layer) and a hydration shell density contrast to bulk water of 0.03 e/Å^3^ as suggested by Svergun et al.^47^ We quantified the deviation between the measured and predicted curves by

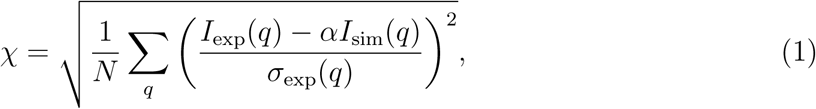

where *α* is a fitted scaling factor (minimizing *χ*) and *σ*_exp_(*q*) is the experimental uncertainty.

Additionally, we compared radii of gyration (*R*_g_) calculated from the IDP simulations and the SAXS experiments. To estimate radii of gyration from the SAXS curves, the Guinier approximation^48^ and the AUTORG software^49^ were used. The error of this estimate (68 % confidence interval (CI)) takes into account propagation of the experimental error and deviations of *R*_g_ between different regions of *q* selected as the Guinier region. To determine *R*_g_ from the simulations, the mean *R*_g_ over all frames was calculated, and errors (68 % CI) were calculated by bootstrapping trajectories.

SAXS curves for MeV-Ntail and for RSP-8 were kindly provided by Longhi et al.^50^ and Rauscher et al.,^1^ respectively. SAXS curves for p53-TAD and CBP-NCBD were measured by Nagy et al.^36^ The SAXS curve for p53-pTAD was also measured by Gabor Nagy (Supp. Methods).

#### 2.3.2 Circular Dichroism Spectra

We compared measured circular dichroism (CD) spectra to those calculated from MD trajectories generated with different force fields. CD spectra were calculated from MD trajectories using SESCA,^51^ which employs a basis set to estimate the CD signal arising from secondary structure elements and individual amino acids. Here, each predicted CD spectrum is obtained as a weighted sum of the basis functions, where the weights reflect the frequency of each secondary structure elements across the simulation trajectories. We used the basis set DS5-4SC1, because it was shown to be accurate for both IDPs and globular proteins.^51^

To quantify the deviation between a predicted and a measured spectrum, both spectra were subsampled in steps of 1 nm in the range of 178-250 nm, because this range is particularly sensitive to secondary structure. The amplitude of the predicted spectrum was then fitted to the measured spectrum with a scaling factor between 0.2 and 5 to account for possible uncertainties in the protein concentration. Using this fitted amplitude, the root mean squared deviations (RMSD) between the subsampled measured and the predicted spectra in thousand mean residue ellipticity units (kMRE) were calculated.

The CD spectra of TIM, HEWL, and RNaseA were obtained from the Protein Circular Dichroism Data Bank (PCDDB),^52^ entries CD0000070000,^53^ CD0000045000,^53^ and CD0000063000,^53^ respectively. The CD spectrum of MeV-Ntail was kindly provided by Longhi and co-workers.^54^ CD spectra of p53-TAD, RSP-8, and CBP-NCBD were measured by Nagy and co-workers.^36^ The CD spectrum of p53-pTAD was also measured by Gabor Nagy (Supp. Methods).

#### 2.3.3 Chemical Shifts

Chemical shifts of C*_α_* atoms were previously measured by NMR and compared to those calculated from our MD trajectories. Chemical shifts were predicted from the MD trajectories using Sparta+.^55^ The difference between measured and predicted chemical shifts was quantified by their RMSD in ppm. This RMSD captures both deviations of the relative frequency of conformational states within the simulated ensembles and the accuracy of the structures of these conformations.

C*_α_* chemical shifts of MeV-Ntail and RSP-8 were kindly provided by Gely et al.^56^ and Rauscher et al.,^1^ respectively. C*_α_* chemical shifts for CBP-NCBD and p53-TAD were obtained from the Biological Magnetic Resonance Data Bank (BMRB),^57^ entries 16 363,^58^ and 17 760,^59^ respectively. A subset of the chemical shifts for p53-TAD was used for p53-pTAD.

#### 2.3.4 Combining Experimental Comparisons of IDP ensembles

To enable a direct comparison between prediction accuracies for different force fields across different experiments, we first normalized the RMSD (for CD and NMR measurements) and *χ* (for SAXS measurements) values obtained from comparing the simulation results with the experimental results. To that aim, for each combination of protein and experiment, e.g., NMR measurements of MeV-Ntail, from each RMSD (or *χ*) value corresponding to each force field its mean value across all force fields was subtracted. Next, we divided each of the resulting values by the standard deviation of all values. To combine the prediction accuracies for SAXS, CD, and NMR experiments of five IDPs into a single number for each force field, we calculated the mean and standard deviation of all normalized RMSD (or *χ*) values obtained for that force field. We refer to this mean as the overall normalized absolute deviation from experiment of a force field.

The statistical significance of accuracy differences between any two force fields was assessed by paired samples *t* -tests. To this end, the normalized RMSD values of both force fields for a specific combination of IDP and experiment were considered a paired sample, of which 15 were used for each comparison (6 for C36m+Tip4p). Every combination of protein and experiment contributes only one RMSD value, but contains multiple uncorrelated data. For a statistical analysis, the total number of uncorrelated data, i.e., degrees of freedom, *n*_dof_ must be estimated as it affects the calculated statistical significance. Here, we assume four degrees of freedom for every combination of protein and experiment and thus a total *n*_dof_ = 60 (24 for C36m+Tip4p). To illustrate, four degrees of freedom of a SAXS curve could correspond to the magnitude of three main axes of an ellipsoidal electron density and a population of transient tertiary structure. The CD spectra computed from our MD trajectories are a weighted sum of six basis spectra corresponding to different (partly transient) secondary structure elements, hence four degrees of freedom is a conservative estimate here. Regarding the NMR comparison, assuming four degrees of freedom corresponds to every fifth C*_α_* chemical shift being independent for the smallest peptide considered here (RSP-8, 20 shifts considered). Overall, we expect *n*_dof_ = 60 to be a conservative estimate, such that the true statistical significance might be higher than the one reported.

#### 2.3.5 Crystal Structures

To assess the stability of the globular proteins nNOS, HEWL, RNaseA, and TIM using the C36m+OPC force field, we compared structures obtained from our simulations to their respective crystal structures. To this end, each snapshot from the MD trajectories was rigid-body fitted to the crystal structure, minimizing the RMSD of all atom positions. This minimal RMSD is particularly sensitive to large conformational changes and partial unfolding of the protein. Specifically, any RMSD *>* 10 Å we observed was due to partial unfolding. The first 100 ns of each trajectory were omitted for this analysis.

#### 2.3.6 Photo-induced Electron Transfer Excitation Decay

The cysteine-tryptophan (C-W) contact-formation kinetics of AGQ-repeat peptides were measured in photo-induced electron transfer (PET) experiments and compared to those computed from our MD trajectories.

In the PET experiments, the tryptophan side chains were excited to an electronic triplet state by a nanosecond ultraviolet light pulse, and the excited state population *p*_exc_(*t*) was monitored over time (Supp. Fig. S9). In the absence of quenchers, the exited state has a slow natural decay time *τ*_0_ of 0.1-1 ms. Upon contact formation with a cysteine residue at a distance *d*_CW_ *≈* 4 Å between cysteine and tryptophan, the excitation is quenched with a much faster quenching time *τ*_q_.^60^ Assuming Markovian two-state contact kinetics with formation time *τ*_+_ and dissociation time *τ_−_* and *τ*_+_ *≫ τ*_q_,^60^ C-W contact-formation kinetics was observed via the decay time

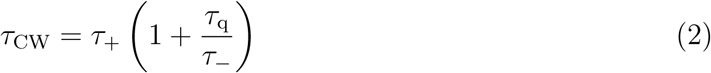

of the measured excited state population *p*_exc_(*t*). In the case of multiple non-contact conformational states or photo-damage, multiple contact formation pathways with possibly different decay times exist, implying a more complex decay of *p*_exc_(*t*). Accordingly, to obtain *τ*_CW_ from the experiment, *p*_exc_(*t*) was described by a convolution *p̃*_exc_(*t*) = *f*_in_(*t*) *∗ f*_out_(*t*) of an effective influx to the excited triplet state with a multiple channel decay. The influx, generated by the laser excitation from the ground state to an excited singlet state and subsequent decay to the triplet state, was described as *f*_in_(*t*) *∝* exp [*−*(*t − t*_on_)*/τ*_in_]Θ(*t − t*_on_), with an onset time *t*_on_, decay constant *τ*_in_, and Heaviside step function Θ; the multi-channel decay was modeled by a sum of exponential functions 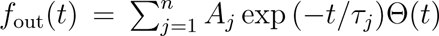 with decay times *τ_j_* and onset at *t* = 0. Put together, for *t > t*_on_,

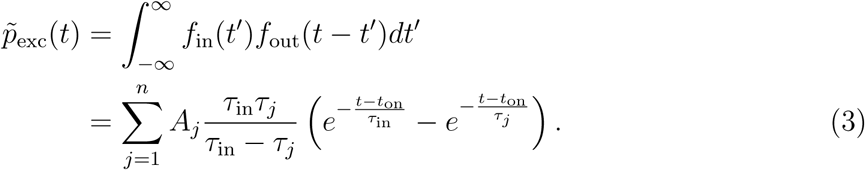

For each peptide, several measurements were carried out and fitted using the same *τ_j_*, *τ*_in_, and *t*_on_, but individual amplitudes *A_j_*. The fastest decay time *τ*_CW,exp_ = min*{τ_j_}*obtained for every peptide was selected for comparison to the *τ*_CW,sim_ calculated from the simulations. The uncertainty of *τ*_CW,exp_ was estimated using the error of the mean of *τ*_CW,exp_ values obtained by fitting to each individual measured curve (Supp. Fig. S10).

Whether the slower decay components are also due to intramolecular C-W contact formation or due to other quenching processes such as interaction with other amino acids, or with buffer molecules, is unclear. We note that our above approach neglects the effect of the natural decay and other possible quenching channels, which might decrease *τ*_CW,exp_. However, we expect this effect to be small, because *τ*_CW,exp_ is faster than *τ*_0_ by over two orders of magnitude and also faster than the second smallest *τ_j_* by more than one order of magnitude (Supp. Tab. S4).

To fit Eq. (3) to *p*_exc_(*t*), parameters ***θ*** = *{t*_on_*, τ*_in_*, τ_j_, A_j_}* were sampled from the Bayesian posterior

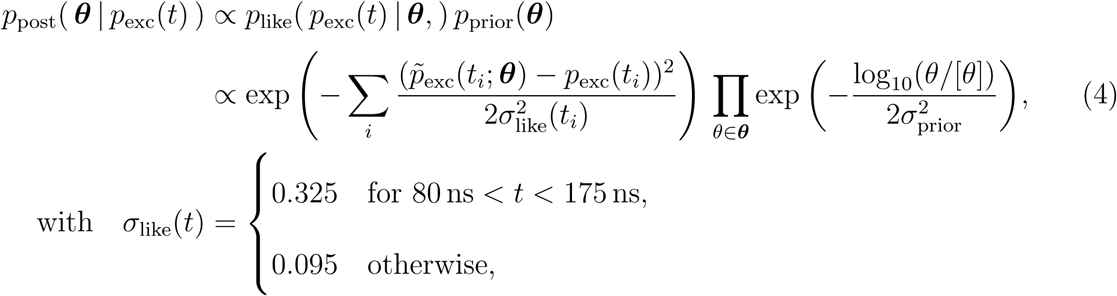

accounting for different uncertainties in high-frequency measurements near the excitation and low-frequency measurements elsewhere (Supp. Fig. S11). A lognormal prior with *σ*_prior_ = 1 was used and [*θ*] denotes the unit of a parameter, which was [*t*_on_] = [*τ*_in_] = [*τ_j_*] = 1 µs, and [*A_j_*] = 1. As a heuristic trade-off between the goodness of the fit and model complexity, the number of exponential components was chosen as *n* = 4, for which the smallest Bayesian information criterion^61^ was obtained for *n* between 2 and 8 and for each of the six peptides (Supp. Fig. S12).

Gibbs sampling was performed for 10^5^ steps at *T* = 1 with adaptive noise and a target acceptance rate *p*_acc_ = 0.5 for each parameter. For each peptide, the set of parameters that minimized Eq. (4) were selected to obtain *τ*_CW,exp_ and are provided in Supp. Tab. S4.

To predict *τ*_CW,sim_ from the MD trajectories, we classified all frames into ‘C-W contact’ and ‘non-contact’ by applying a cut-off distance of 4 Å to the smallest distance *d*_CW_ of any atom of the tryptophan indole to the cysteine sulfur atom. From the obtained time series, the mean contact formation time *τ*_+_ and dissociation time *τ_−_* were calculated. To calculate *τ*_+_, from every non-contact frame, the time until the next contact frame was recorded and their average was calculated; and likewise for *τ_−_*. From these values, *τ*_CW,sim_ was calculated using eq. (2). Because the required *τ*_q_ cannot be obtained from the simulations, nor were we able to measure it, we resorted to previous measurements, ^60,62,63^ which reported values between 1 ns and 5 ns. To account for this uncertainty, we estimated *τ*_q_ independently for each force field by fitting (minimizing *χ*^2^) predicted *τ*_+_ and *τ_−_* to the experimentally determined *τ*_CW,exp_ using eq. (2), including both experimental errors and errors from trajectory-wise bootstrapping.

To rank the accuracy of different force fields, their average accuracy for six different peptides, S-(AGQ)5-C-(AGC)1,3,5-W and C-(AGQ)5-S-(AGC)1,3,5-W, was compared. To quantify this accuracy, two different scores were used. First, the mean absolute Z-score 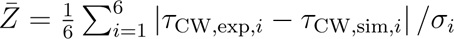 captures the deviation between the measured relaxation times τCW,exp,i and the predicted ones τCW,sim,i for all peptides i in units of their respective standard errors. These errors 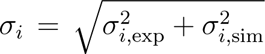 take into account measurement errors and uncertainty due to limited sampling of the simulations. The latter was calculated by bootstrapping trajectories. The error of *Z̅* was estimated by bootstrapping the six *Z*-values. Second, the 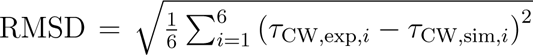 captures the absolute deviation between predicted and measured relaxation times. Its error was calculated using Gaussian error propagation.

The experimental setup of the pump-probe apparatus was identical to previous work by Otteson et al.^22^ Peptides were synthesized by Tufts University, with reported purities ranging from 85 % to 98 %. Peptide purity was verified by Quantum Time of Flight (QTOF) mass spectrometry. Peptides with *<* 94 % purity were further purified by High-Performance Liquid Chromatography on a C18 semi-prep column to achieve *≥* 94 % purity, then lyophilized and reanalyzed by QTOF prior to use. PET experiments were carried out in 100 mM sodium phosphate buffer pH 6.0 at 20 *^◦^*C using a peptide concentration of 200 µM (*ɛ* = 5500 M*^−^*^1^ cm*^−^*^1^). To reduce dissolved oxygen, samples were bubbled with USP grade nitrous oxide for one hour. This minimizes quenching of the tryptophan triplet state by O_2_, while N_2_O acts as an electron scavenger for electrons generated by water decomposition under the UV laser pulses.

#### 2.3.7 Temperature-jump Infrared Spectroscopy for CLN025

T-jump IR spectroscopy data for CLN025 showed that, at 300 K, its folding behaviour can be described by a two-state model.^32^ Using Van’t Hoff analysis, the thermodynamic parameters of folding Δ*H*_f_ = *−*37.0*±*1.5 kJ mol*^−^*^1^ and Δ*S*_f_ = *−*96.8*±*5.1 J mol*^−^*^1^ K*^−^*^1^ were determined,^32^ yielding a free-energy difference of folding at 300 K of Δ*G*_f_ = *−*8.0 *±* 2.3 kJ mol*^−^*^1^ and a relative population of folded states *p*_f_ = 1*/*[1 + exp(Δ*G*_f_ */k*_B_*T*)] = 96.1 % (68 % CI: 90.8-98.5 %). The folding time at 300 K, *τ*_f_ = 165 *±* 25 ns, was interpolated from a fit of Arrhenius law to measured relaxation times.^32^ The unfolding time *τ*_u_ = 4.1 µs (68 % CI: 1.5-10.3 µs) was calculated using transition state theory as *τ*_u_ = *τ*_f_ exp(*−*Δ*G*_f_)*/k*_B_*T*).

To predict the CLN025 folding kinetics from our simulations, trajectory frames were assigned to either the folded or unfolded states by visual inspection of time traces of the RMSD to the crystal structure and *R*_g_. In cases were assignments were unclear based on these traces, e.g., when *R*_g_ rose slightly higher than usual in the folded state for a short time, structures of the corresponding frames were visualized using PyMol.^64^ Two-state Markov models were fit to the discretized trajectories, using a lag time of *τ*_MSM_ = 10 ps (1 frame) and enforcing detailed balance. To estimate the uncertainty, 20 000 samples were drawn from the Bayesian posterior probability distribution using Markov chain Monte Carlo as implemented in Pyemma2.^65,66^ A log-uniform prior probability distribution was assumed for the transition probabilities.

## 3 Results and Discussion

### 3.1 Intrinsically Disordered Protein Ensembles

To investigate whether MD simulations with the new force field combination C36m+OPC show improved accuracy for IDPs compared to simulations using C36m+Tip3p, we first compared predicted radii of gyration (*R*_g_) to SAXS measurements for five IDPs. For these IDPs, C36m+Tip3p systematically underestimated *R*_g_ values, specifically for larger IDPs (Fig. 1). In contrast, C36m+OPC did not show any systematic deviation and predicted the *R*_g_ of the large MeV-Ntail more accurately.

**Figure 1:**
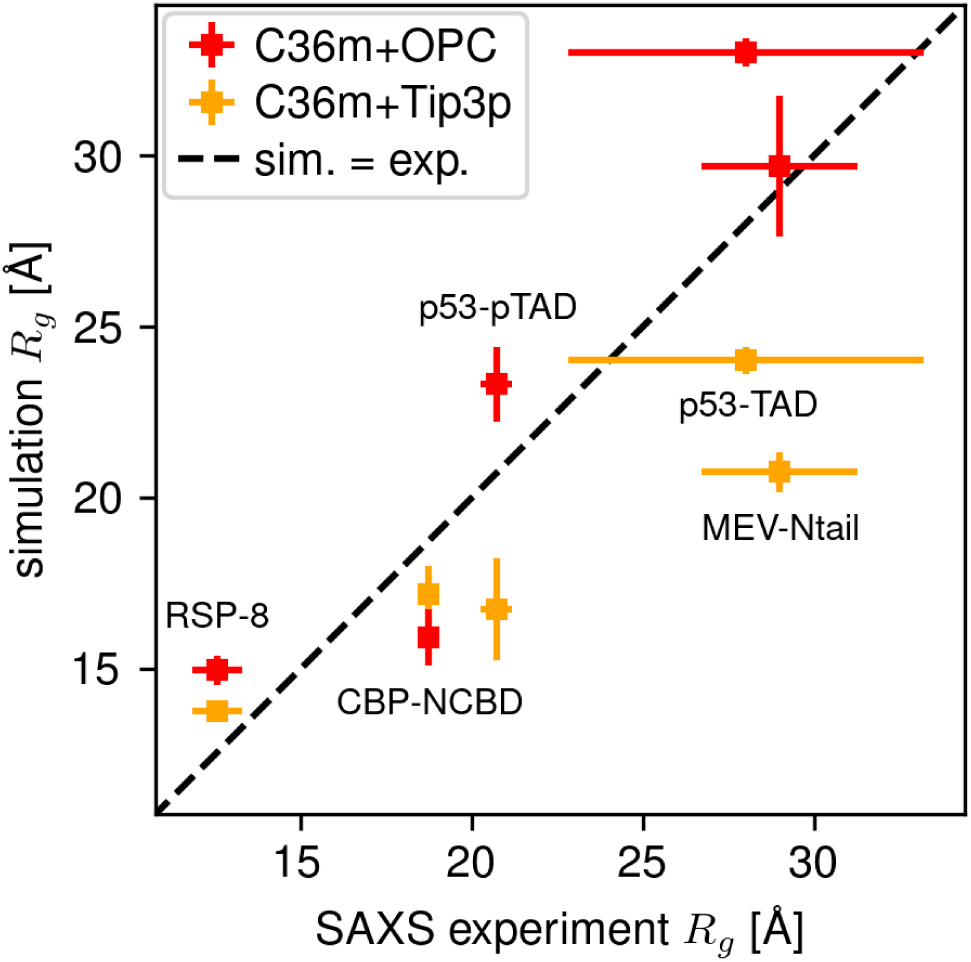
IDP ensemble compactness calculated from simulations compared to experiment. Radii of gyration (*R*_g_) of five IDPs calculated using C36m+OPC (red) and C36m+Tip3p (orange) versus *R*_g_ determined by SAXS experiments. For the calculated values, error bars were estimated by bootstrapping trajectories and represent 68 % confidence intervals. Experimental error bars also represent 68 % CI and take into account propagation of measurement error and uncertainty due to selecting the Guinier region.

This result is a marked improvement over C36m+Tip3p, which raises the question whether other observables that characterize IDP ensembles might also be described more accurately by this force field combination. To assess such overall accuracy, we directly compared predicted SAXS curves, CD spectra, and NMR C*_α_* chemical shifts with measurements.

To this end, we tested not only C36m+OPC and C36m+Tip3p, but also several other force fields (Fig. 2). Specifically, we assessed C36m+Tip4p to see whether changing to a 4-point water model other than OPC has a similar effect. To assess the improvement over older generation Charmm force fields, we also tested C22*. Finally, to estimate how C36m+OPC compares to recent Amber force fields, we tested A03ws, A99disp, and A99sb-ws.

**Figure 2:**
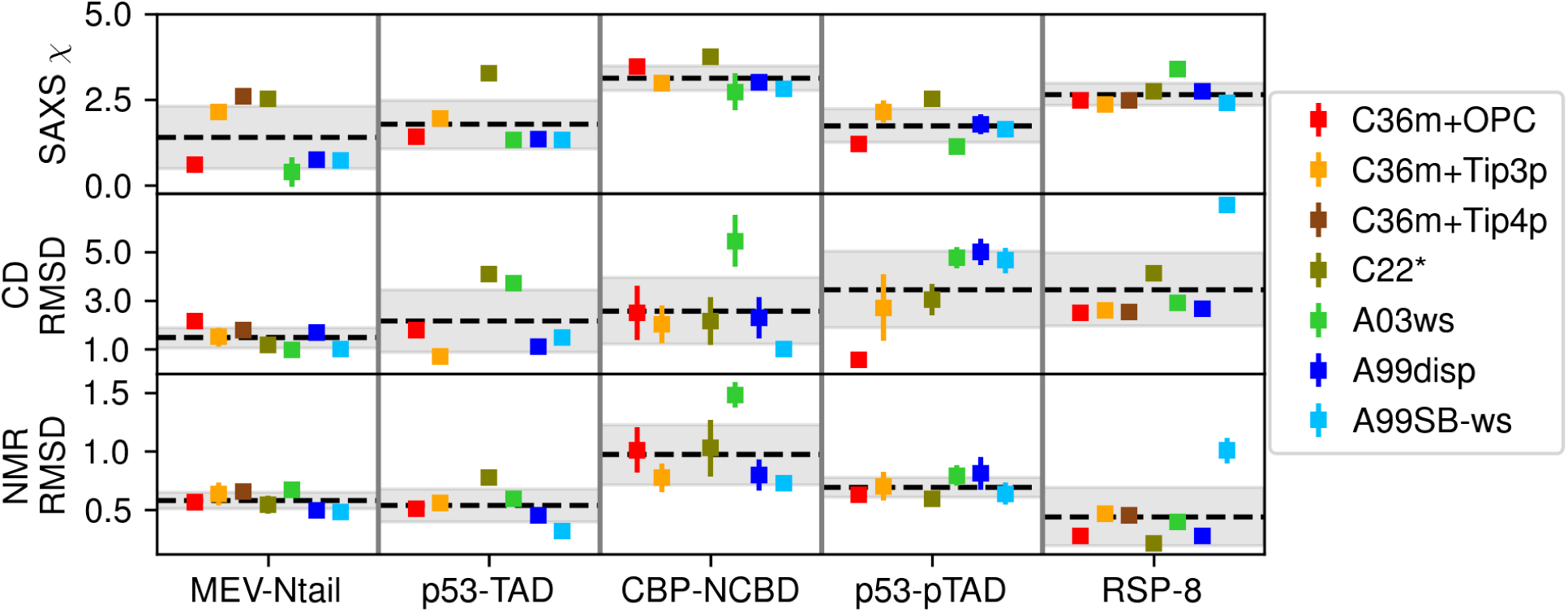
Accuracy of seven force field combinations for IDP ensembles. Using seven different force field combinations, deviations from measures SAXS curves (top row), CD spectra (mid row), and NMR C*_α_* chemical shifts (bottom row) are quantified. Shown are *χ* value after fitting calculated to measured SAXS curves with an overall intensity scaling factor and RMSD values between calculated and measured CD spectra and chemical shifts, respectively. C36m+Tip4p has only been assessed for MEV-Ntail and RSP-8. Error bars were estimated by bootstrapping trajectories and represent 68 % confidence intervals; data points without error bars have errors smaller than the symbol size. Means and standard deviations of the *χ* and RMSD values are shown as dashed black lines and grays shaded areas, respectively.

Figure 2 summarizes our assessments using RMSD-type deviations, as defined in Methods, with lower numbers indicating higher accuracy. Each panel shows the comparison of the tested force fields to a specific experiment of one IDP. As an example, the SAXS measurement of MEV-Ntail (upper left) is more accurately predicted by C36m+OPC and all Amber force fields than by C36m+Tip3p, C36m+Tip4p, and C22*. A similar ranking is obtained for p53-TAD and p53-pTAD, whereas larger deviations from experiment are seen for CBP-NCBD and RSP-8 for all force field combinations. For the CD measurements, a mixed picture is obtained, with high accuracy throughout all force field combinations for MEV-Ntail, good accuracy for some of the force field combinations for p53-TAD, CBP-NCBD, and p53-pTAD, and less accurate results throughout for RSP-8. Mixed results were also seen for the NMR measurements, where, too, all force field combinations performed well for MEV-Ntail and, with few exceptions, for p53-TAD, p53-pTAD, and RSP-8, whereas lower accuracy is achieved throughout for CBP-NCBD. Overall, different rankings are seen, but despite some apparent trends, no obvious patterns emerge.

Towards a more systematic overall assessment, we therefore combined all 15 RMSD values (6 for C36m+Tip4p) into a single number (see Methods), the mean normalized absolute deviation from experiment for each force field (Fig. 3A). To this end, we normalized the RMSDs for each test case by subtracting their mean (dashed black lines in Fig. 2) and dividing by their standard deviations (gray shaded areas in Fig. 2). As can now be seen more clearly in Fig. 3A, A99sb-ws and C36m+OPC show the smallest overall deviation, followed by A99disp and C36m+Tip3p. The remaining force fields C36m+Tip4p, C22*, and A03ws appear to be markedly less accurate. Fig. 3B shows the *p*-values for comparisons of the overall accuracy of any pair of force fields. As can be seen, C36m+OPC is more accurate than C36m+Tip3p (*p <* 0.05), C36m+Tip4p (*p <* 0.01), C22* (*p <* 0.01), and A03ws (*p <* 0.01), but no significant difference was observed to A99disp and to A99sb-ws.

**Figure 3:**
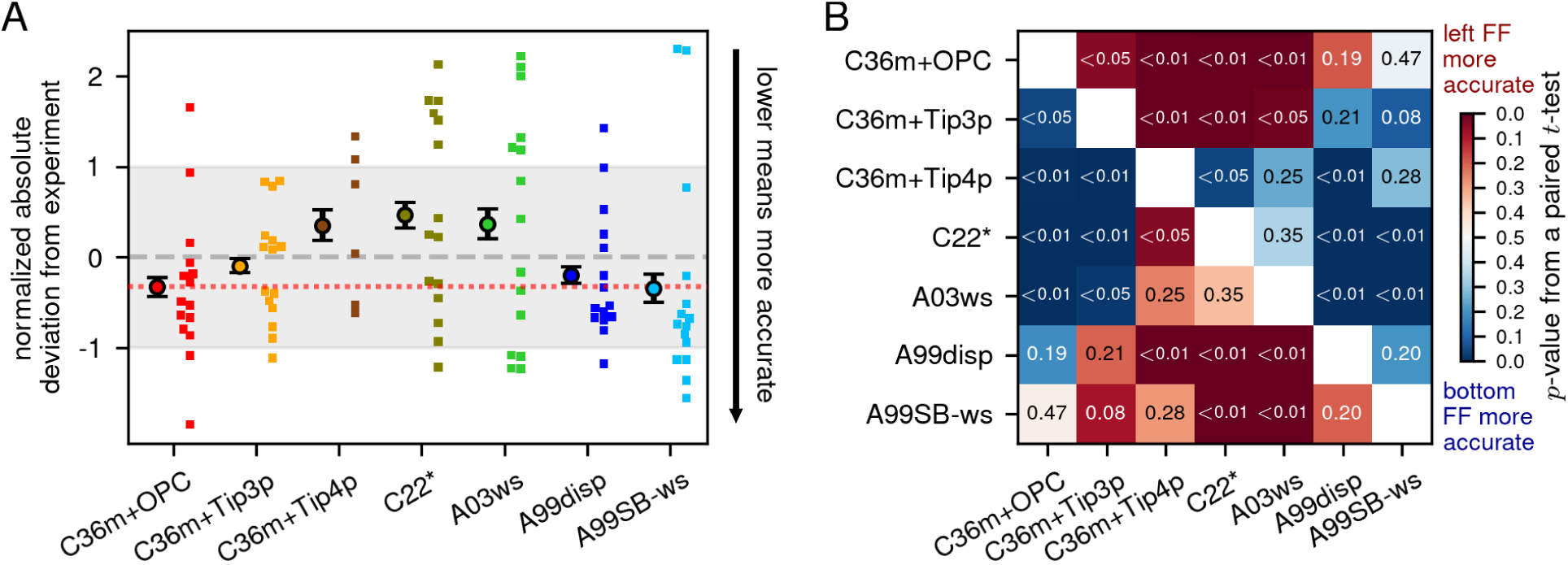
Overall accuracies of IDP ensembles for seven force field combinations. **(A)** Averaged (circles) and individual normalized deviations (small squares) of all observables shown in Fig. 2; deviations were normalized by subtracting means and dividing by respective standard deviations. To aid comparison, the dotted red line indicates the mean deviation of C36m+OPC. Error bars denote the error of the mean assuming four independent degrees of freedom per experiment and per protein, and thus were computed from the standard deviations *σ* as *σ*/√60. They do not capture covariances of paired data; therefore, the statistical significance of differences between the force fields can not be visually inferred from these error bars. **(B)** For comparison of pairs of force fields, the colors and numbers indicate *p*-values for the accuracy differences obtained for all force field pairs.

The gain in accuracy of C36m+OPC over C36m+Tip3p may be attributable to the switch from a 3-point to a 4-point water model. To test this hypothesis, we additionally tested a combination of C36m with a different 4-point water model, namely C36m+Tip4p, for MeV-Ntail and RSP-8. Because the accuracy of C36m+Tip4p was lower than that of C36m+Tip3p (*p <* 0.01) and C36m+OPC (*p <* 0.01), we conclude that the observed gain in accuracy of C36m+OPC over C36m+Tip3p is not primarily due to a generally higher accuracy of 4-point water models.

The used normalization scheme is just one possible way to combine RMSD values into a single score and might introduce a bias. To check for such a possible bias, we tested several other schemes, such as calculating mean and standard deviations of the RMSD values for each experiment across all proteins instead of for individual proteins (Supp. Fig. S1), scaling the deviations to different experiments without subtracting the mean (Supp. Fig. S2), or applying a threshold for good or acceptable fits to experiments (Supp. Fig. S3). These other combination methods rank C36m+Tip4p as more accurate than C22* and A03ws, and A99sb-ws as slightly less accurate than C36m+OPC. The sensitivity of the rank of C36m+Tip4p to the combination method is likely due to its small sample size of only two proteins. Overall, we found no marked effect of the choice of assessment scheme on the outcome.

In summary, our assessment shows that for the tested IDPs C36m+OPC achieves the accuracy of the best performing Amber force fields (A99sb-ws and A99disp). This is particularly noteworthy in light of the fact that the OPC water model was not designed specifically for use with Charmm36m (nor the other way around), whereas A99disp was developed together with its specific water model to achieve high accuracy for IDPs and is thus widely regarded as one of the most accurate protein force fields.^2,15,22,67^

### 3.2 Globular Proteins

The accuracy gain of C36m+OPC compared to C36m+Tip3p for IDPs might come at the expense of decreased accuracy for globular proteins. In particular, the stronger protein-water interactions that helped C36m+OPC favor less compact IDP ensembles may also destabilize globular proteins or perturb their ensembles. To test if this is the case, we also assessed force field accuracy on several folded proteins by comparing to NMR C*_α_* chemical shifts, CD spectra, and crystal structures.

With respect to C*_α_* chemical shifts (Fig. 4A), C36m+OPC (red), C36m+Tip3p (orange), and A99sb-ildn (purple) show similar accuracy. A99disp (blue) is slightly more accurate than other force fields for proteins XPCB, nNOS, and HEWL. Also the CD spectra calculated from all tested force fields show similar accuracy (Fig. 4B), except HEWL and TIM, for which A99disp and A99sb-ildn, respectively, are slightly more accurate.

**Figure 4:**
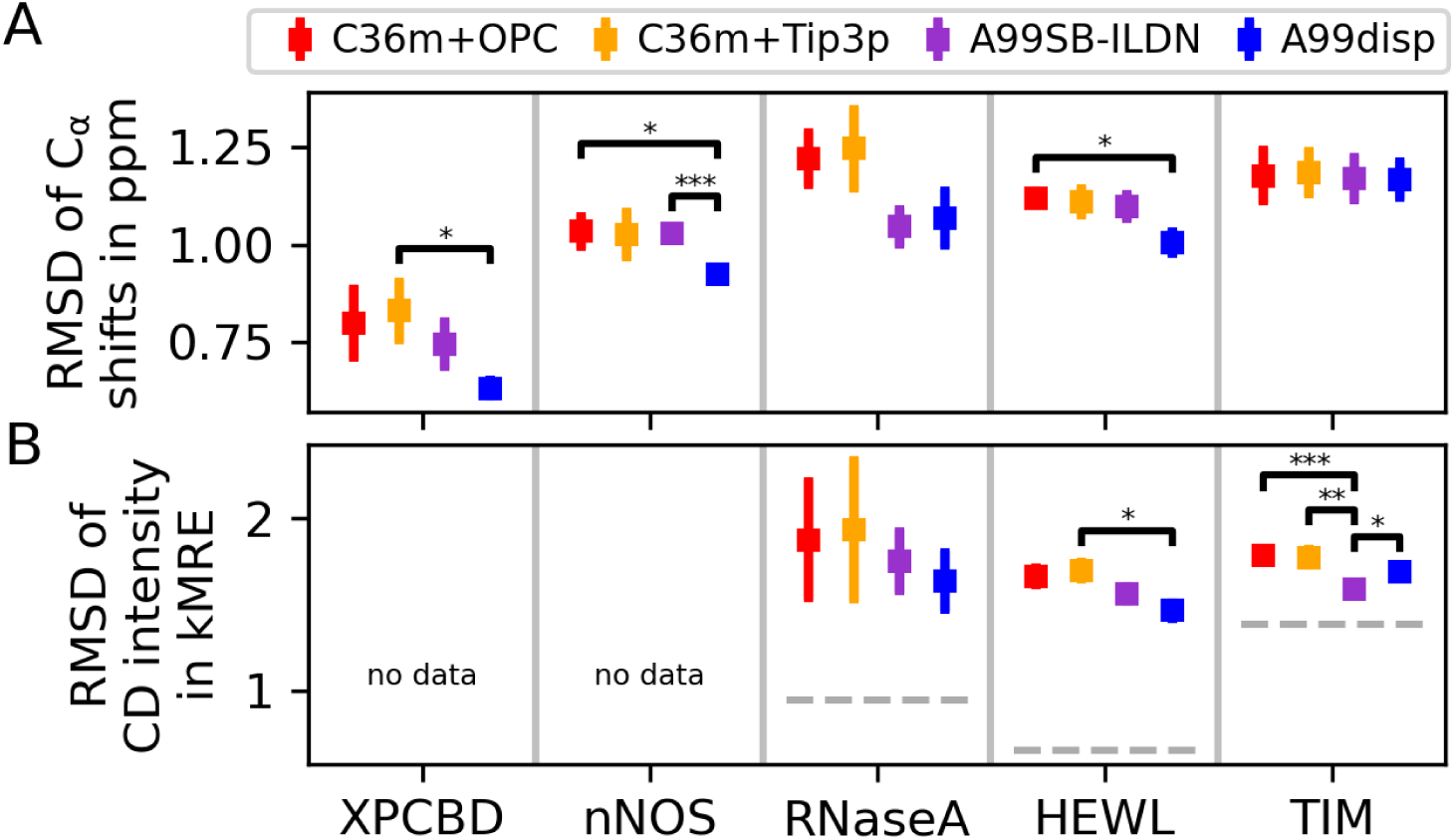
Overall accuracies for several globular proteins for four force field combinations. (**A**) RMSD between C*_α_* chemical shifts measured in NMR experiments and predicted from simulations with different force fields (colors). (**B**) RMSD between measured and predicted CD spectra for three different globular proteins. The dashed gray lines indicate the best possible prediction given the used basis set. Error bars were estimated by bootstrapping trajectories and represent 68 % confidence intervals. All significant differences are marked using *p*-value star notation.

To assess how close the trajectories remained to the crystal structures of globular proteins using C36m+OPC or C36m+Tip3p, Fig. 5 shows time traces of the RMSD between atom positions in simulations and in the crystal structures. Partial unfolding (RMSD *>* 10 Å) is observed for C36m+Tip3p for RNaseA in 5/20 trajectories, for TIM in 1/20 trajectories, and for nNOS in 1/50 trajectories; and using A99disp for nNOS in 11/50 trajectories. In contrast, no unfolding is seen for C36m+OPC and A99sb-ildn. This observation indicates either that globular proteins are more stable using C36m+OPC than using C36m+Tip3p, or at least that unfolding kinetics are slower in C36m+OPC, or both. Besides partial unfolding events, the RMSD traces of C36m+OPC and C36m+Tip3p are similar, as are those of A99sb-ildn and A99disp. Compared to the Charmm force fields, the Amber force fields show smaller mean RMSD values for HEWL, RNaseA, and TIM, and larger mean RMSD values for nNOS.

**Figure 5:**
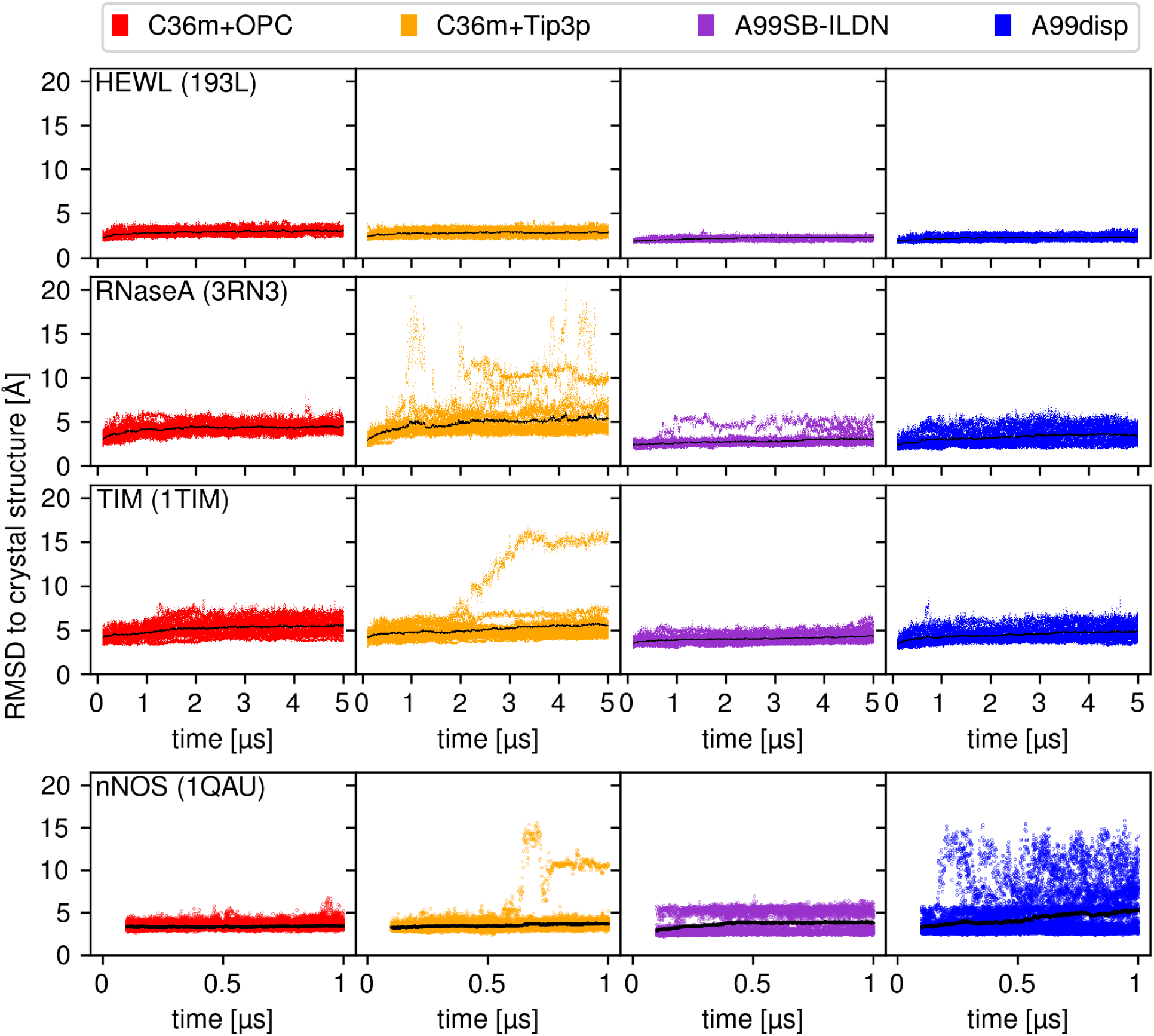
Deviations of globular protein structures relative to their crystal structures. Each panel shows superimposed time traces of the RMSD of C*_α_* atom positions from the respective crystal structure (PDB ID in brackets) for 20 simulations (colored dots; 50 for nNOS) and their mean (black line) for four globular protein (rows) and four force fields (columns, colors).

Overall, all tested force fields show similar accuracy for globular proteins. Specifically, C36m+OPC does not seem less accurate than C36m+Tip3p. We neither found evidence for C36m+OPC destabilizing globular proteins compared to C36m+Tip3p, nor for decreased accuracy of their ensembles. These results suggest that the increase in accuracy for IDPs in C36m+OPC did not come at the cost of decreased accuracy for globular proteins.

### 3.3 AGQ-repeat Peptide Kinetics

To also assess how accurately C36m+OPC predicts kinetic properties, we compared cysteine-tryptophan (C-W) contact dynamics predicted from simulations of the intrinsically disordered S-(AGQ)_5_-C-(AGQ)_1,3,5_-W and C-(AGQ)_5_-S-(AGQ)_1,3,5_-W peptides to relaxation times measured by photo-induced electron transfer (PET) experiments. In these PET experiments, the population of excited-state tryptophan indoles is monitored over time after photo-excitation. When an excited tryptophan indole forms a van der Waals contact with a cysteine sulfur atom, the excited state is quenched via electron transfer much faster than the unperturbed decay, with a quenching time constant *τ*_q_. Hence, the measured population after excitation decays with an effective decay time *τ*_CW_, which reports on the kinetics of contact formation.

To calculate *τ*_CW_ from our simulations, we extracted the C-W contact formation times *τ*_+_ and dissociation times *τ_−_* for each peptide and force field (Supp. Tab. S5), and fitted the peptide-independent quenching time *τ*_q_ to the measured *τ*_CW_ using Eq. (2) (see Methods). For all force fields, the obtained quenching rates lie within the range 1 ns *≤ τ*_q_ *≤* 5 ns (Table 1) which agrees with previous measurements. ^60,62,63^

**Table 1:** Fitted PET quenching times *τ*_q_; *Z̅* and RMSD values compare AGQ peptide simulations with four different force field combinations to experiments.

| force field | $\tau_q$ [ns] | $\bar{Z}$ | RMSD [ns] |
| --- | --- | --- | --- |
| C36m+OPC | $5.0 \pm 0.9$ | $1.6 \pm 0.4$ | $800 \pm 500$ |
| C36m+TIP3P | $5 \pm 10$ | $11 \pm 7$ | $320 \pm 120$ |
| A03ws | $3.5 \pm 0.4$ | $3.3 \pm 1.3$ | $78 \pm 23$ |
| A99disp | $1.80 \pm 0.25$ | $2.1 \pm 0.8$ | $200 \pm 40$ |

Figures 6A,B compare calculated and measured relaxation times due to PET *τ*_CW_ as a function of C-W sequence separation. Most calculated *τ*_CW_ correlate well with the measured ones. The most pronounced deviations are that C36m+OPC significantly overestimates *τ*_CW_ for C-W separation 15 and that C36m+Tip3p and A99disp underestimate *τ*_CW_ for the end-to-end contacts at C-W separations 19, 25, and 31. As a side note, for A03ws the obtained *τ*_q_ = 3.5*±*0.4 ns deviates markedly from the value *τ*_q_ = 1.25 ns previously calculated from simulations using the same force field for similar peptides.^62^

**Figure 6:**
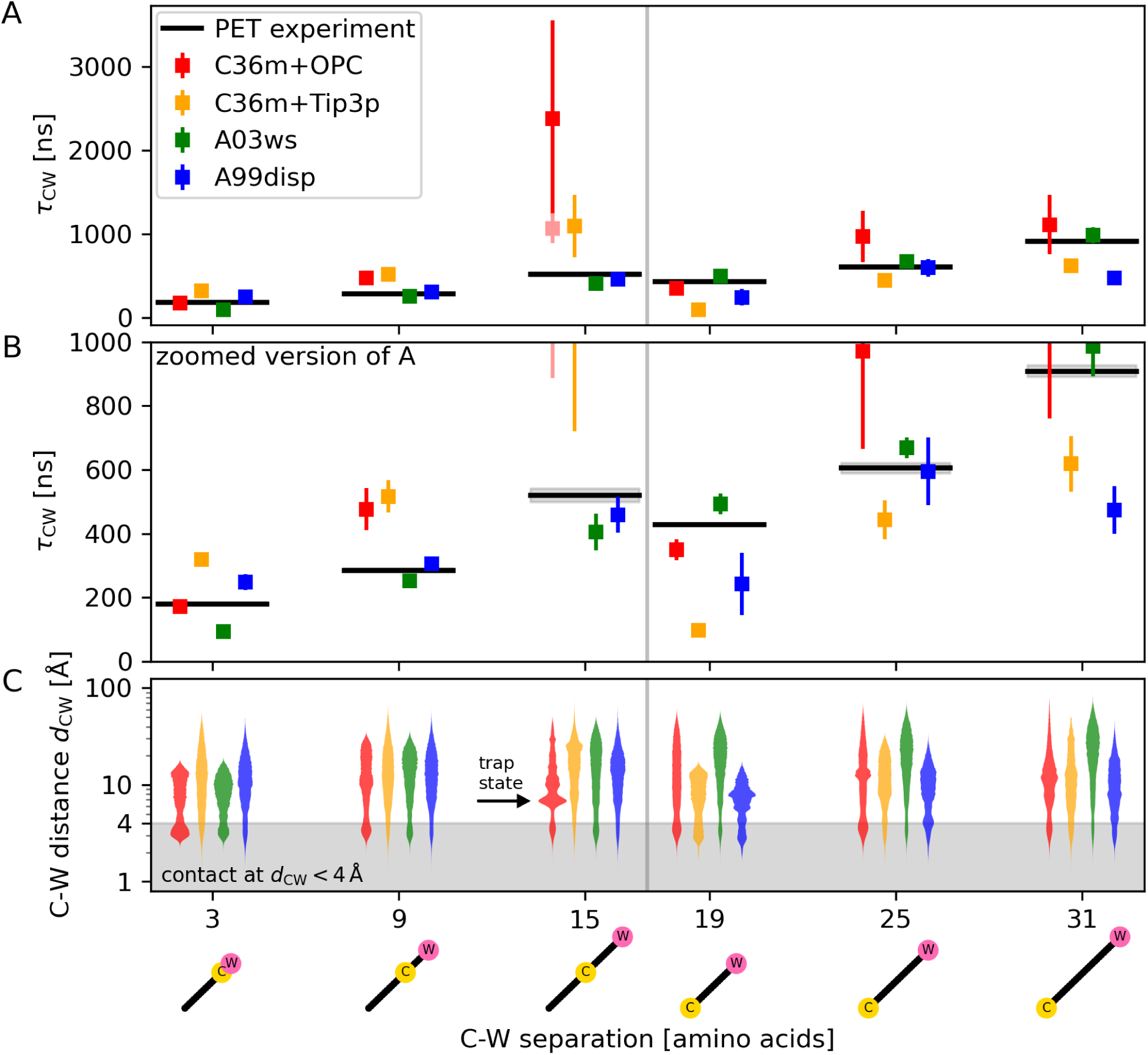
Calculated C-W contact kinetics of AGQ-repeat peptides compared to PET measurements. (**A**) For six different peptides, relaxation times due to PET *τ*_CW_ were calculated from simulations using different force fields (colors) and are shown for different C-W sequence separations (*x*-axis). For each force field combination, *τ*_q_ was obtained from fits of the calculated to the measured relaxation times (black) using eq. (2). Error bars of calculated *τ*_CW,sim_ were estimated by bootstrapping trajectories. Error bars and measurement errors (gray area) represent 68 % confidence intervals. Points are slightly horizontally displaced for visual clarity. (**B**) Vertically expanded view of panel A. (**C**) Distributions of C-W distances *d*_CW_ obtained for each peptide and force field (colors); the gray area indicates the contact region, *d*_CW_ *<* 4 Å (gray area). The pictograms below indicate C and W positions in the peptide sequence.

Table 1 quantifies the overall accuracy for each force field by their mean *Z*-score *Z̅* and the RMSD between calculated and experimental values. Based on the RMSDs, A03ws reproduces the kinetics more accurately than A99disp, which in turn is more accurate than C36m+Tip3p. The ranking of C36m+OPC is unclear due to large uncertainty and requires closer analysis.

Notably, the large RMSD of C36m+OPC is due to a single trajectory at C-W separation of 15 amino acids, which corresponds to the end-to-center contact in the long peptide S-(AGQ)_5_-C-(AGQ)_5_-W, whose formation time is overestimated (Fig. 6A). These slow kinetics are caused by a kinetic trap state at *d*_CW_ *≈* 0.7 nm (Fig. 6C, arrow), which prevents contact formation in one of five simulations (Supp. Fig. S8). This ‘anecdotal sampling’ is reflected in the large uncertainty shown Fig. 6A, which results in only marginal statistical significance of this outlier. This finding is also supported by a low *Z̅* = 1.6 *±* 0.4, which implies that for C36m+OPC the deviations from the experiment are largely due to limited sampling and do not necessarily point to larger force field inaccuracies. Indeed, if the trapped state was a rare event, one would expect that better agreement is achieved for an analysis omitting the trajectory with the trapped state, which is actually what we observed (light red color symbol in Figs. 6A,B).

In contrast, C36m+Tip3p deviations show a markedly higher significances, with *Z̅* = 11 *±* 7, implying that these are mainly not due to limited sampling. Rather, C36m+Tip3p systematically underestimates the formation time of end-to-end contacts (C-W separation of 19–31 amino acids) compared to end-to-center contacts (C-W separation of 3–15 amino acids). Because the deviations for C36m+Tip3p are both systematic and statistically significant, they point to force field inaccuracies. One possible reason could be that the Tip3p* water model overestimates the self-diffusion coefficient, as has been reported previously.^68^

Taken together, and despite the limited statistics, our results suggest that AGQ-peptide kinetics are more accurately described by C36m+OPC than by C36m+Tip3p.

### 3.4 CLN025 Folding Kinetics

As a second, independent benchmark for protein kinetics, we compared predicted and measured folding and unfolding rates of the synthetic 10-residue miniprotein CLN025 at 300 K. Figure 7 compares rates calculated from simulations using C36m+OPC, C36m+Tip3p, A99sb-ildn, and A99disp with measurements by equilibrium and T-jump IR spectroscopy experiments.^32^ The T-jump measurements suggest that CLN025 folding follows a two-state process at 300 K.^32^ Based on the measured equilibration rates,^32^ we estimated a folding time of 150 ns (68 % CI: [100, 200]) and from the measured thermodynamic parameters we calculated a fraction of folded states of 0.961 (68 % CI: [0.878, 0.988]). From these numbers we estimated an unfolding time of 3.6 µs (68 % CI: [0.7 µs, 16.4 µs]).

**Figure 7:**
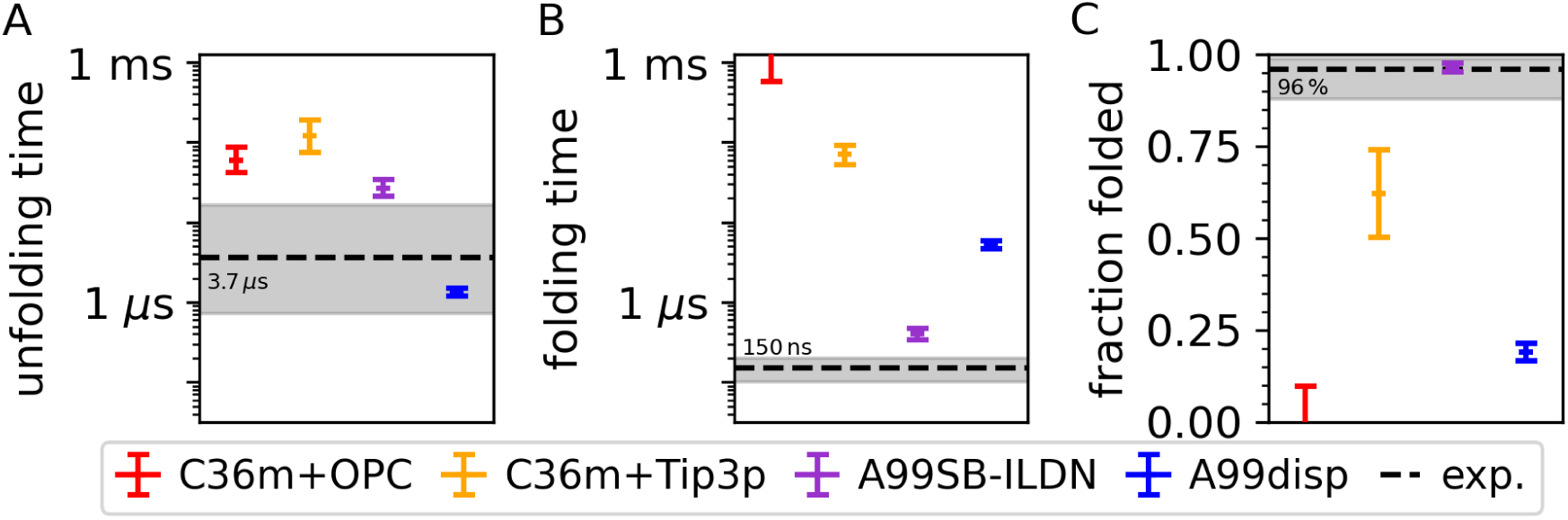
Calculated folding kinetics of CLN025 compared to experiment. Four different force field combinations have been used (colors) to calculate (**A**) mean unfolding times, (**B**) mean folding times, and (**C**) the fraction of folded states from the simulations. Equilibrium and T-jump measurements are shown as dashed lines, gray areas indicate 68 % confidence intervals. Because no folding events were observed in the C36m+OPC simulations, only lower and upper limits for the folding time and the relative folded population, respectively, are shown.

As a prerequisite for the analysis, we examined the simulated conformational ensembles. All force fields predicted metastable folded and unfolded states (Supp. Fig. S15). Additionally, C36m+OPC, C36m+Tip3p, and A99disp predicted a metastable molten globule state, and A99disp and A99sb-ildn predicted a metastable misfolded state. A slight variation of the misfolded state including an additional hydrogen bond between Thr6 and Trp9 side chains was observed using A99sb-ildn. The other intra-protein hydrogen bonds in the misfolded state closely resemble those of the folded state. If the misfolded state exists *in-vitro*, this close resemblance would probably render it indistinguishable from the folded state in IR spectroscopy. In contrast, the molten globule state observed in simulations lacks all hydrogen bonds probed in the IR experiment^32^ and therefore would be indistinguishable from the unfolded state. Accordingly, for comparison with experiments, the misfolded and molten globule states observed in the simulation were treated as part of the folded and unfolded state populations, respectively.

Only the unfolding time calculated from the A99disp simulations agrees with experiment, all other force fields overestimate it (Fig. 7A). Also the folding time is overestimated by all tested force fields (Fig. 7B), with A99sb-ildn being closest to the experiment, followed by A99disp and C36m+Tip3p. No folding event was observed using C36m+OPC, such that only a lower limit for the folding time is shown. Figure 7C shows the fraction of folded states calculated from folding and unfolding times. Only the folded fraction calculated using A99sb-ildn agrees with experiment, all other force fields underestimate it. Contrary to experiment, C36m+OPC and A99disp predict faster unfolding than folding, leading to folded fractions below 50 %.^31,32^

Taken together, A99sb-ildn predicts CLN025 folding kinetics most accurately. It predicts the folded fraction accurately and overestimates folding and unfolding times by a factor of 2.7 and 7.4, respectively. C36m+Tip3p predicts the kinetics too slow by two orders of magnitude and the folded fraction agrees with experiment only qualitatively (faster folding than unfolding). Using C36m+OPC, we observed unfolding but no folding event, hence the predicted kinetics qualitatively disagree with experiment and are clearly slower than in C36m+Tip3p. Therefore, combining C36m with OPC decreased its accuracy of CLN025 folding kinetics relative to C36m+Tip3p.

## 4 Conclusions

In this study, we assessed the accuracy of a new combination of force field and water model C36m+OPC^19^ and compared its accuracy to the accuracy of established force fields, including the original combination C36m+Tip3p^4^ and A99disp,^2^ which is widely considered one of the most accurate protein force fields. To this end, we performed extensive MD simulations of a number of peptides and proteins, calculated SAXS curves, CD spectra, and NMR C*_α_* chemical shifts, and compared these to measurements. These systems include five IDPs with diverse sizes, transient secondary structure propensities, and transient tertiary structures. In addition to these ensemble based observables, we assessed the accuracy of the force fields with respect to peptide kinetics. In particular, we compared calculated relaxation times due to PET with measured ones for a set of disordered AGQ-repeat peptides with diverse sizes, and also compared predicted folding and unfolding times of CLN025 to T-jump IR spectroscopy experiments. To further check if the new force field combination compromises the accuracy of simulations of globular proteins, we compared CD spectra and C*_α_* chemical shifts predicted from simulations of three and five globular proteins, respectively, with respective measurements. Finally, to asses globular protein stability, we compared structural changes during our simulations to crystal structures.

Our main finding is that combining C36m with the OPC^19^ water model tends to increase simulation accuracy compared to the canonical C36m+Tip3p for IDP ensembles for most observables, without compromising accuracy of globular proteins. Importantly, C36m+OPC avoids the systematic underestimation of the radius of gyration of IDP ensembles observed for C36m+Tip3p. Additionally, C36m+OPC predicts SAXS curves, CD spectra, and C*_α_* chemical shifts for the five tested disordered proteins more accurately than C36m+Tip3p and most other tested force fields. For all tested globular proteins, C36m+OPC and C36m+Tip3p predicted similarly accurate CD spectra and C*_α_* chemical shifts, indicating that the increased protein-water interactions (or possibly inconsistent protein and water parameters) of C36m+OPC do not markedly perturb globular protein ensembles. Furthermore, C36m+OPC maintains the stable crystal structures of globular proteins, as does C36m+Tip3p in most replicas. Overall, we think that the increased accuracy of C36m+OPC outweighs the increased computational cost of moving from a 3-point to a 4-point water model, especially for IDP simulations.

Our second main result is that C36m+OPC shows similar accuracy for IDP ensembles to A99sb-ws and A99disp, which is generally considered highly accurate for IDPs and globular proteins,^2,15,30^ superseding most other tested force fields such as C36m+Tip3p, C22*, and A03ws. C36m+OPC and A99disp also show similar accuracy for globular proteins, indicating that C36m+OPC reaches state-of-the-art accuracy for general protein force fields.

We speculate that the increase in accuracy of C36m+OPC over C36m+Tip3p for IDPs is due to the more accurate balance between protein-protein, protein-solvent, and solvent-solvent interactions interactions of the OPC water model, which was optimized for electrostatics. One aspect of this success is that OPC is a 4-point water model, while Tip3p is a 3-point water model. The mere fact of using a 4-point water model, however, does not automatically yield higher accuracy, as demonstrated by the less accurate C36m+Tip4p combination.

As a third result, our kinetics benchmarks suggest that, whereas C36m+OPC reproduces relaxation times due to PET of AGQ-repeat peptides more accurately than C36m+Tip3p, it fails to accurately predict kinetic rates of CLN025 folding. While no tested force field completely agrees with the measured CLN025 folding kinetics, A99disp accurately predicts the unfolding time, A99sb-ildn accurately predicts the folded fraction, and C36m+Tip3p at least qualitatively captures faster folding than unfolding. From these mixed results we can not draw general conclusions on the accuracy of C36m+OPC for kinetics beyond the peptides tested here.

Further improvements to C36m+OPC are likely possible by adjusting protein force field parameters, as was done for A99disp or in the work of Gong et al.^15^ More generally, based on our and other recent results,^22,23^ we consider conformational and folding kinetics of proteins an important target for force field evaluation and optimization, which so far received too little emphasis.

## Acknowledgement

N.K. thanks the International Max Planck Research School for Physics of Biological and Complex Systems for financial support. We thank Reinhardt Klement for providing the simulations of p53-TAD using A03ws and Paul Robustelli for providing the simulations of MeV-Ntail using A99disp. We thank Sarah Rauscher for providing the simulations using C22*, the SAXS curve, and the NMR chemical shifts of RSP-8.

## Author Contributions

G.N. performed simulations of the IDPs and AGQ-repeat peptides. G.N. and N.K. analyzed the simulations of the IDPs and AGQ-repeat peptides. N.K. performed and analyzed simulations of the globular proteins and CLN025. G.N. and H.G. initiated and conceptualized the project. C.F.O. performed the PET experiments. C.F.O. and N.K. analyzed the PET experiments. S.M.V. supervised the PET experiments. N.K. created the figures and wrote the manuscript. L.V.B., H.G. and S.M.V. revised the manuscript. H.G. supervised the project.

## Supporting Information Available

- Three different other combination methods for the RMSD/*χ* values shown in Fig. 2
- Measured and predicted SAXS curves, CD spectra, and C*_α_* chemical shifts
- C-W distance traces from simulations of S-(AGQ)_5_-C-(AGQ)_5_-W using C36m+OPC
- Measured time traces of excited triplet state tryptophan populations from PET experiments
- *τ*_CW_ from fits to single PET curves used for error estimation
- Visualization of regions with different experimental uncertainty in the PET curves
- Bayesian information criterion plots for choosing the number of exponential components to fit PET curves
- Analysis of the effect of choosing different cutoff distances for C-W contact formation on the calculated PET decay times
- Visualization of CLN025 free energy landscape and selected representative structures
- RMSD to the crystal structure and *R*_g_ of CLN025 trajectories
- Molecular Dynamics simulation parameters
- Best-fit parameters for measured PET relaxation curves
- Calculated C-W contact formation and dissociation times for all force fields
- Description of SAXS and CD experiments of p53-pTAD
- Description of starting structures for IDP and AGQ-peptide simulations

## TOC Graphic

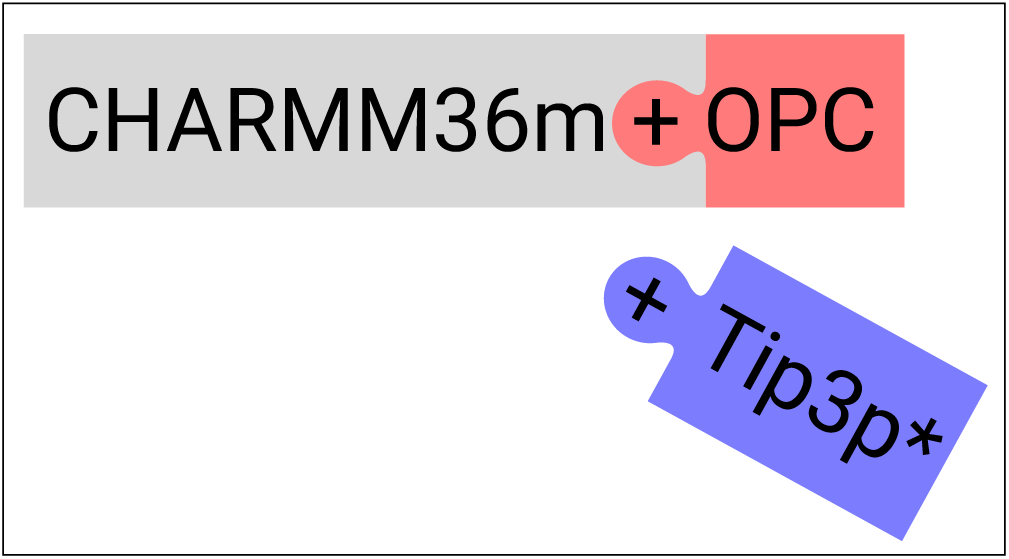

