## Supplementary Material for "Combining CHARMM36m with OPC water improves accuracy"

#### Supplementary Figures and Tables

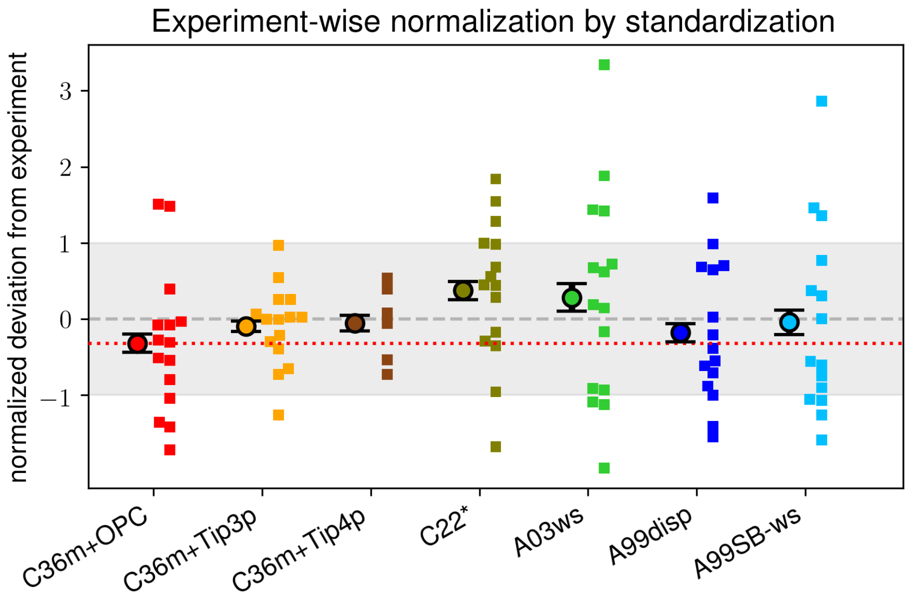


Figure 1: **Overall accuracies of IDP ensembles for seven force field combinations based on experiment-wise normalization.** Averaged (circles) and individual normalized deviations (small squares) of all observables shown in Fig. 2. Here, RMSD and $\chi$ values were normalized as in the main analysis, separately for each experiment, but considering all proteins.

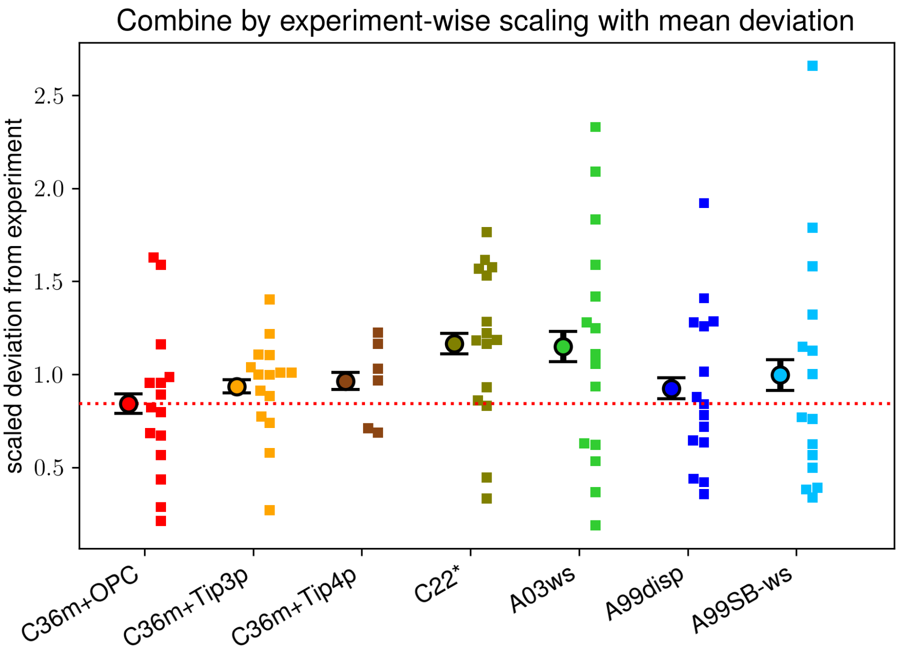


Figure 2: **Overall accuracies of IDP ensembles for seven force field combinations based on experiment-wise scaling.** Averaged (circles) and individual normalized deviations (small squares) of all observables shown in Fig. 2. Here, RMSD and $\chi$ values were normalized by dividing by the experiment-wise mean RMSD across all proteins and force fields. This scaling made RMSD magnitudes comparable across experiments, while preserving the original mean (i.e., without mean-centering beforehand).

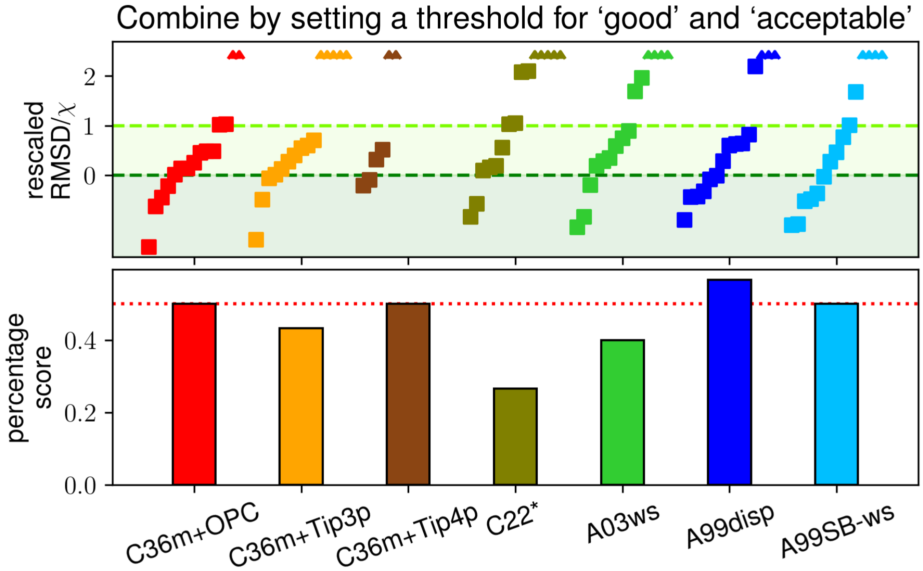


Figure 3: **Overall accuracies of IDP ensembles for seven force field combinations based on categorizing predictions.** Force field accuracy was categorized into good and acceptable predictions. The upper panel shows for each force field, all RMSD and $\chi$ values that were normalized by subtracting the threshold for good accuracy and subsequently dividing by the difference between the thresholds of good and acceptable accuracy (compare Fig. X). For SAXS $\chi$, the thresholds for good and acceptable were 1.0 and 2.0, respectively. The corresponding thresholds were 2.0 and 3.0 for CD RMSD and 0.5 and 1.0 for NMR $C_\alpha$ chemical shift RMSD. The lower panel shows the percentage of achievable score (good accuracy in all cases), with acceptable accuracy earning half the score of good accuracy.

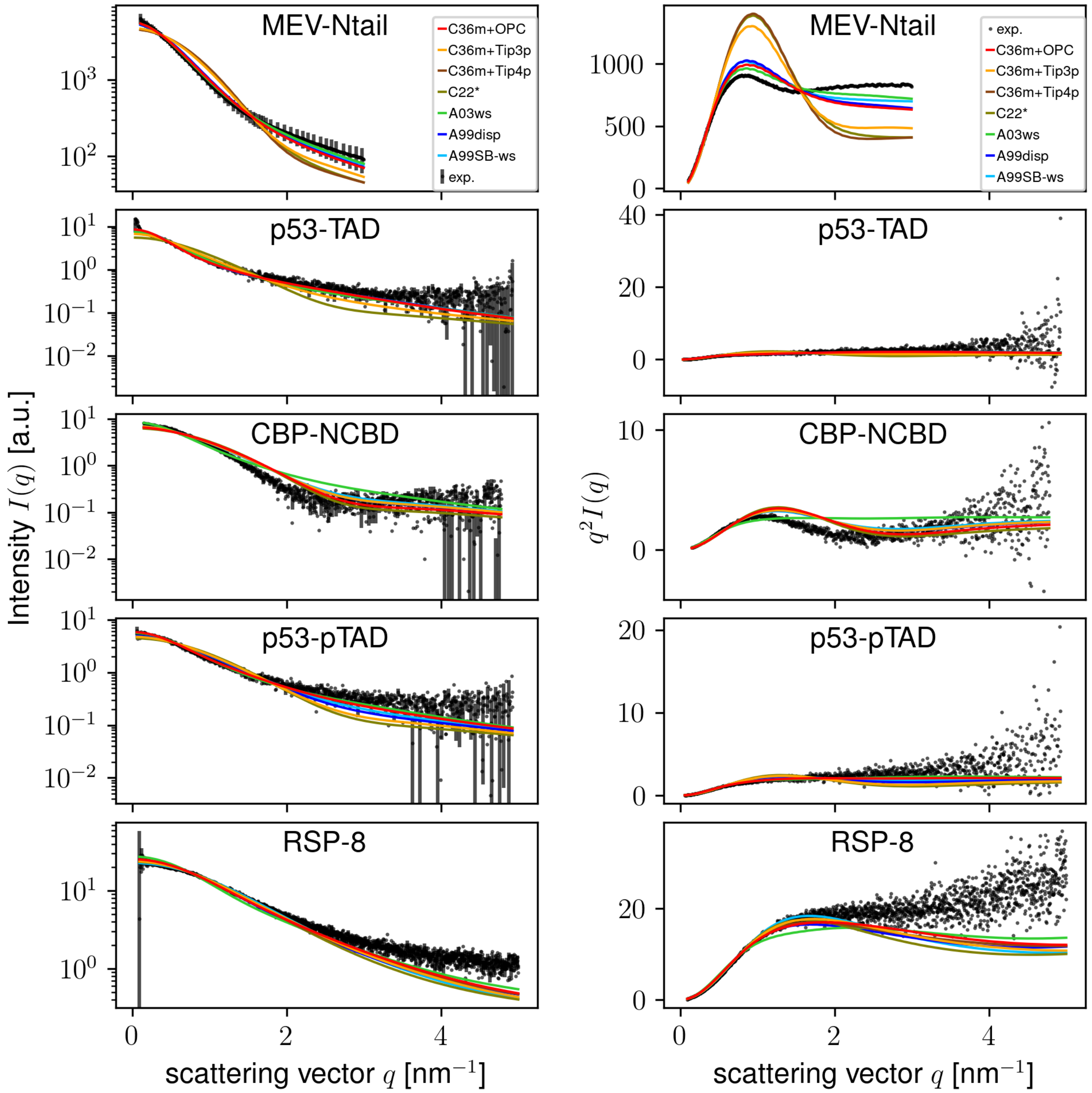


Figure 4: **Comparison of measured and calculated SAXS curves.** Raw curves and Kratky plots are shown in the left and right column, respectively. Calculated curves were scaled to fit the measured one.

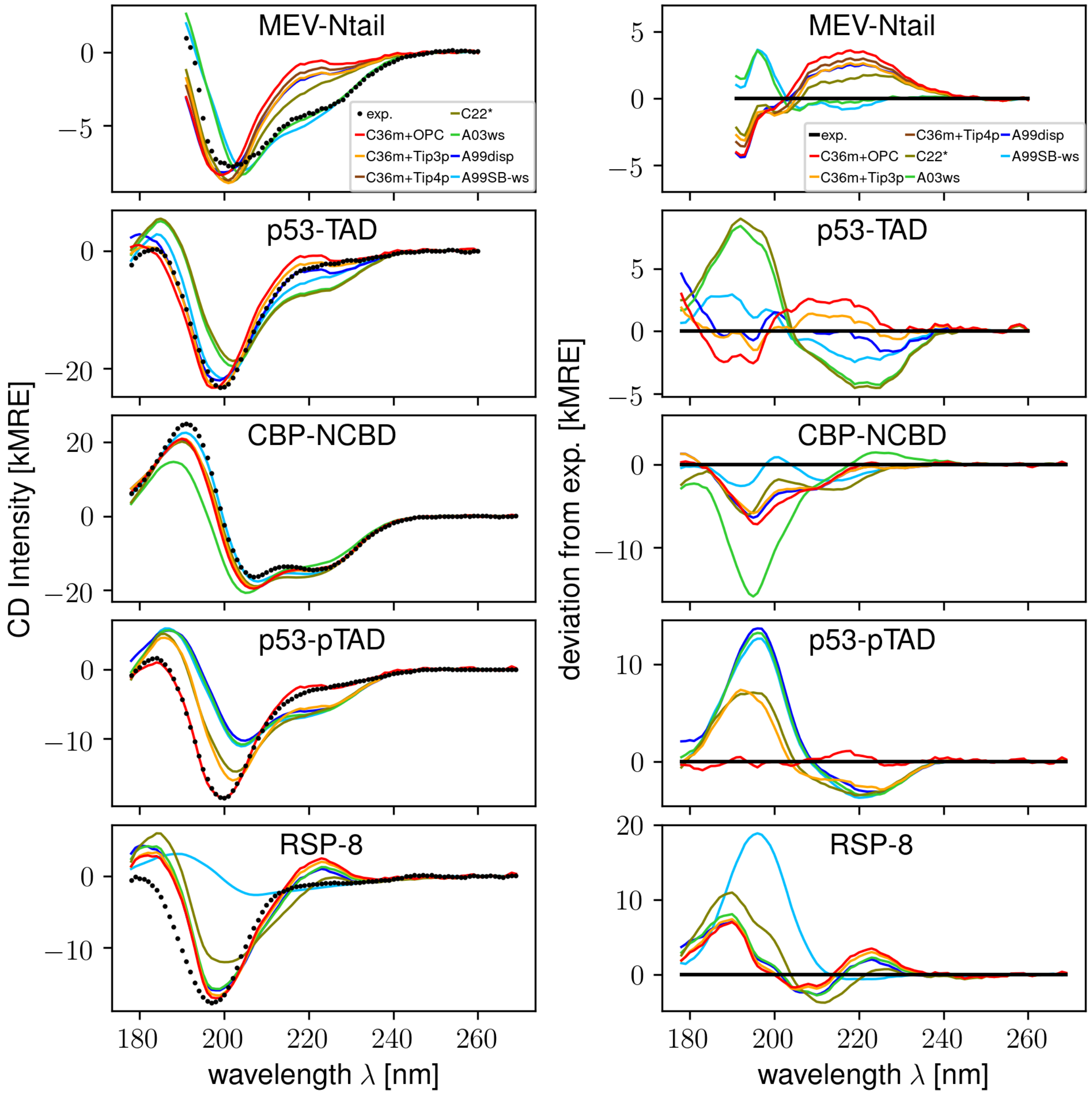


Figure 5: **Comparison of measured and calculated CD spectra.** Raw spectra are shown in the left column. The right column shows the deviation between calculated and measured spectra. Calculated spectra were scaled to match the measured spectrum (Methods).

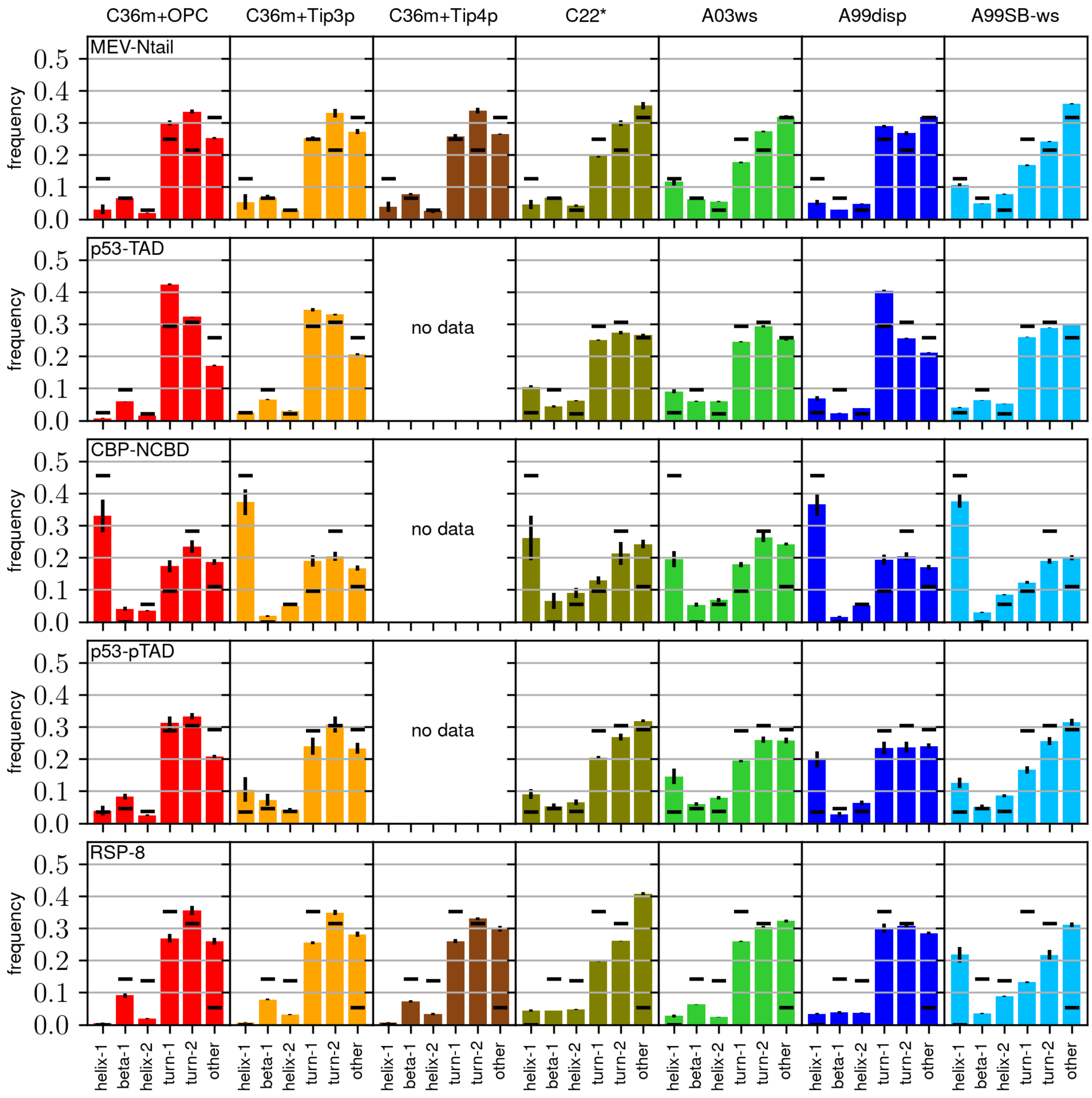


Figure 6: **Weights for CD basis spectra from different force fields (columns, colors).** Weights are calculated from predicted secondary structure content for each protein (rows). Black horizontal lines represents optimal weights from fitting the basis set to the measured spectra.

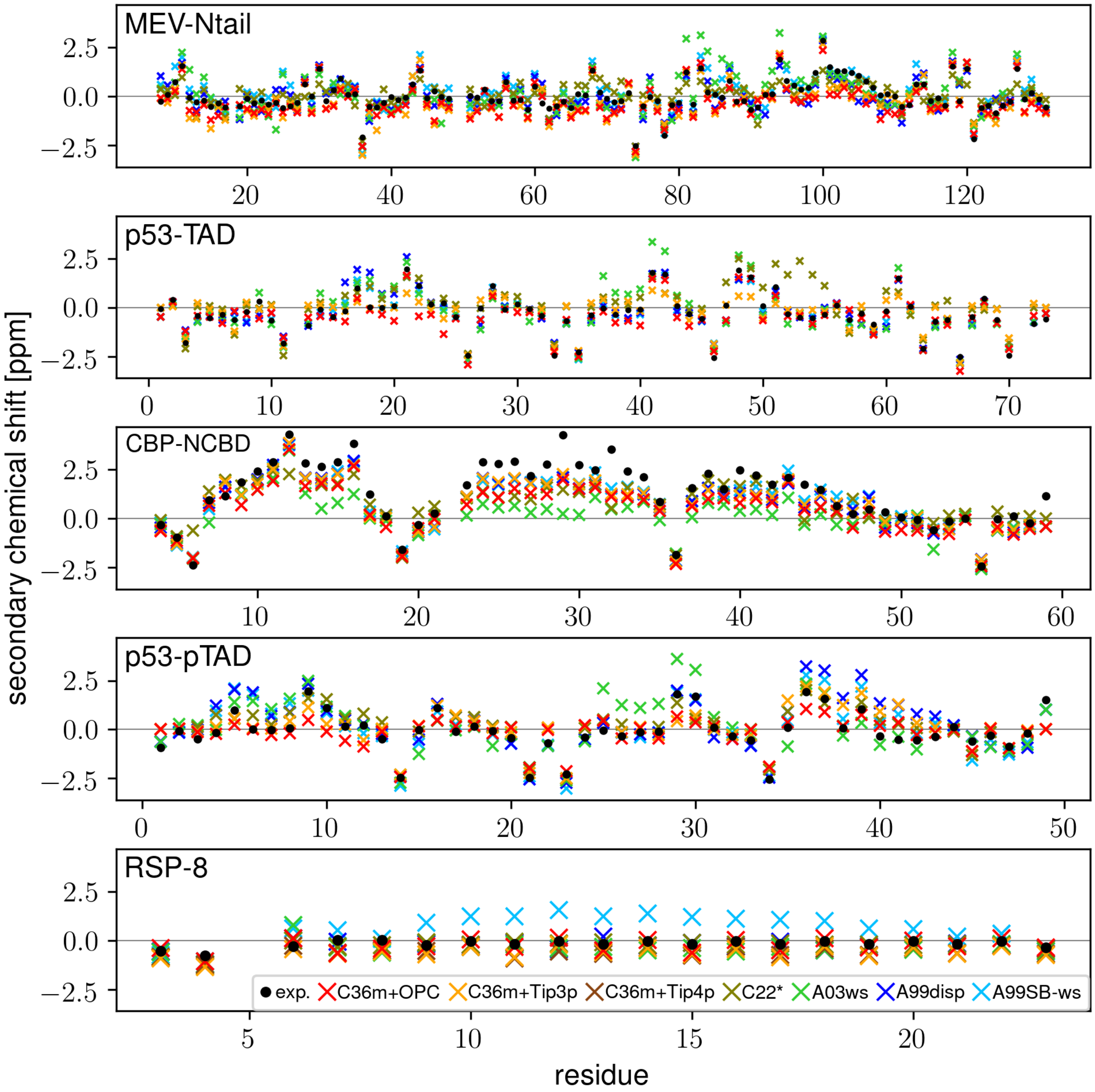


Figure 7: **Comparison of measured and calculated $C_\alpha$ chemical shifts.**

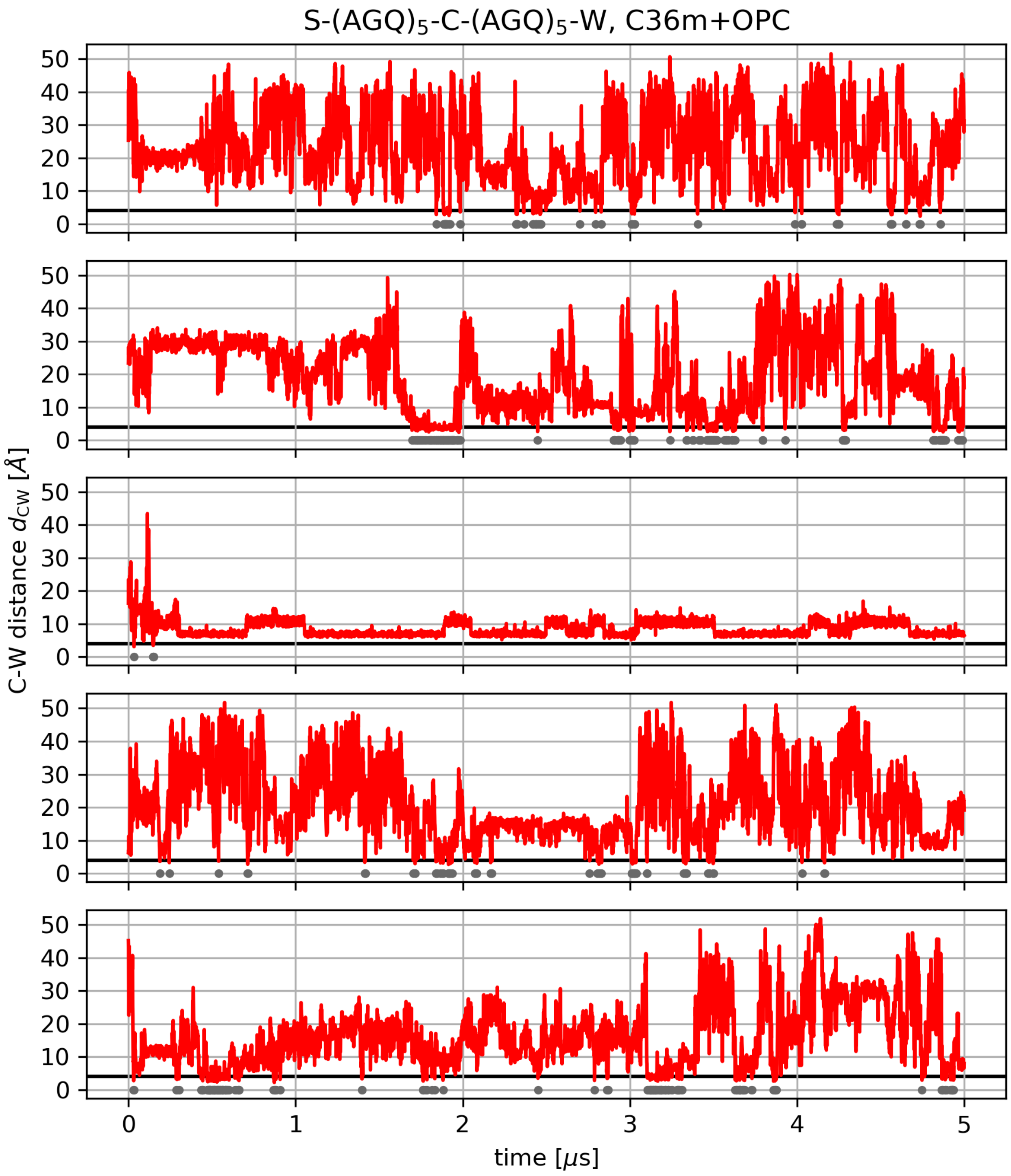


Figure 8: **C-W contact dynamics from MD simulations.** Time traces of the C-W distance $d_{\mathrm{CW}}$ (red) from simulations of S-(AGQ)$_5$-C-(AGQ)$_5$-W using C36m+OPC. The black line marks the threshold for contact formation at 4 Å. Contact frames are marked with gray circles.

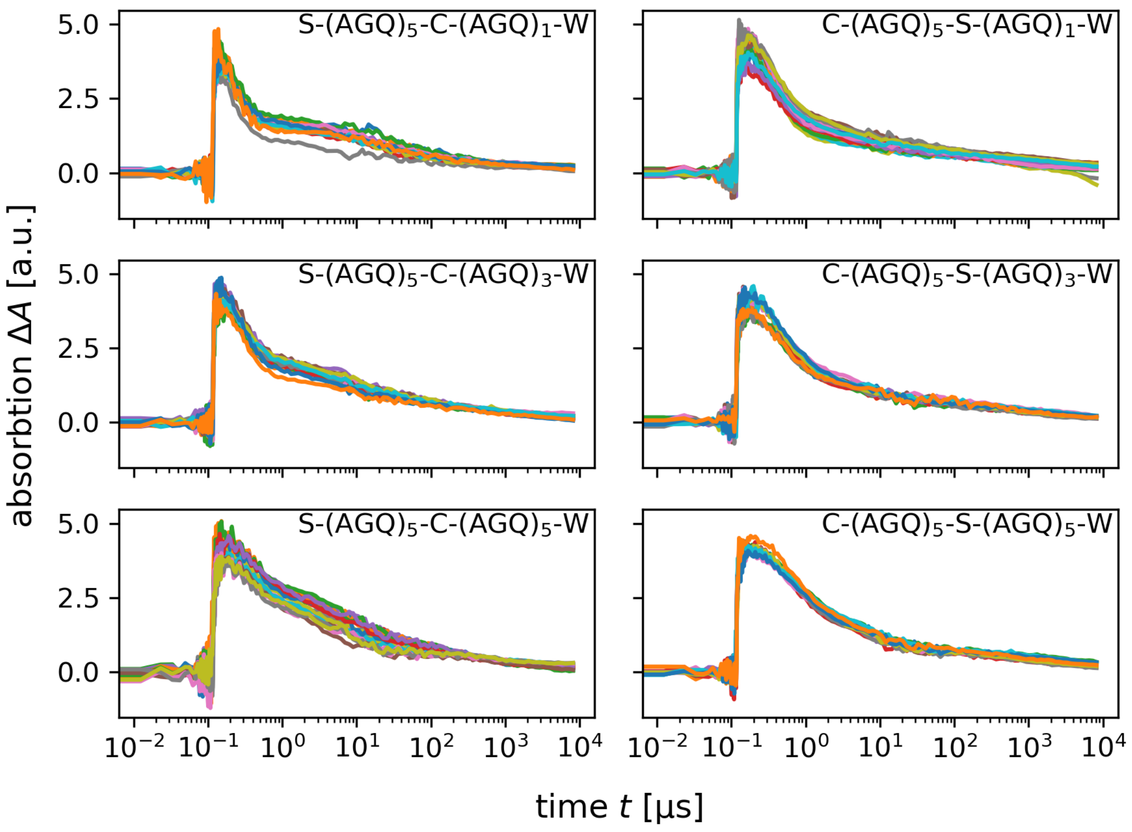


Figure 9: **Time traces of the absorption $\Delta A$ measured at $458\,\mathrm{nm}$ in PET experiments of six AGQ-repeat peptides.** The measured absorption is indicative of the population of excited triplet state tryptophan $p_{\mathrm{exc}}$. For each peptide, 12 independent time traces were measured (colors; 19 for S-(AGQ)$_5$-C-(AGQ)$_5$-W, 20 for C-(AGQ)$_5$-S-(AGQ)$_1$-W).

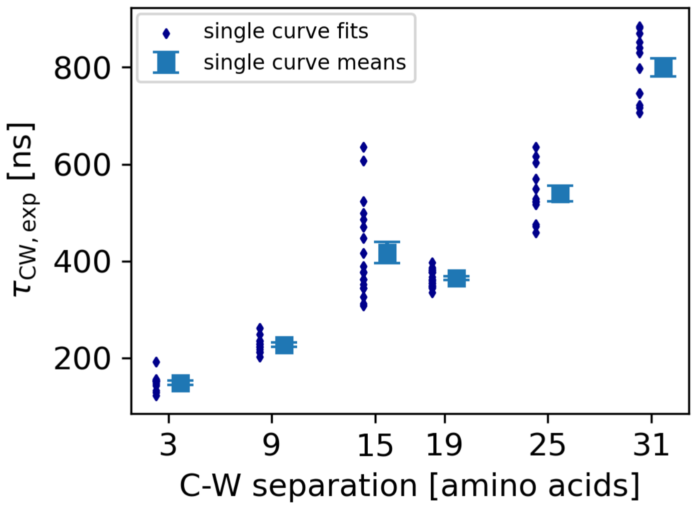


Figure 10: **Decay times due to PET over C-W separation estimated from single-curve fits.** Decay times $\tau_{\mathrm{CW,exp}}$ obtained from individual curves are shown as blue diamonds and their means as blue squares. Error bars represent the error of the mean.

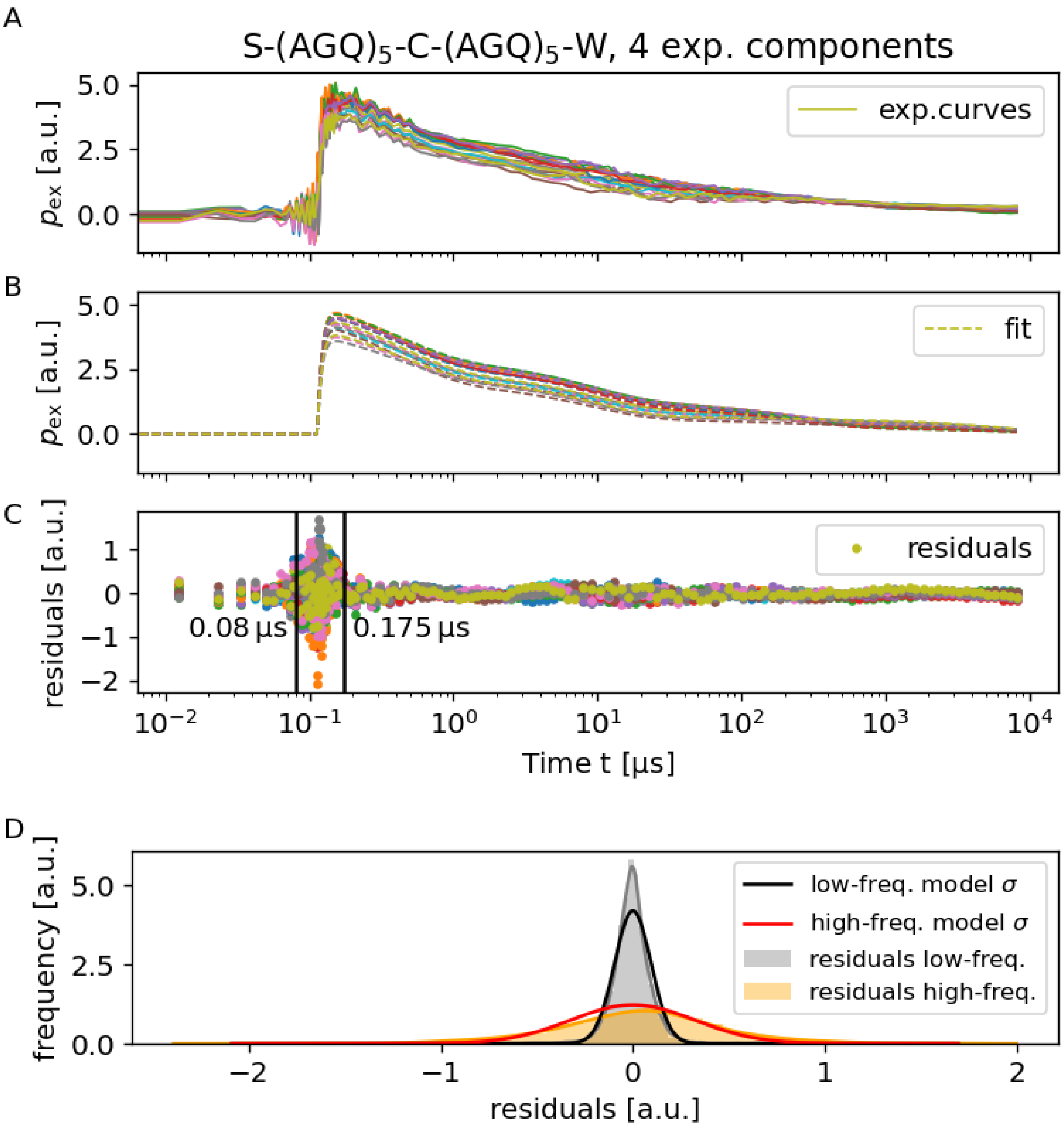


Figure 11: **Fluctuations in PET measurements differed markedly in the high-frequency and low-frequency regime, requiring a two-region error model for fitting.** (**A**) Measured triplet state population of a selected peptide (S-(AGQ)$_5$-C-(AGQ)$_5$-W) as a function of time (solid lines) and global fit thereof with $n = 4$ exponential components (dashed lines). (**B**) Difference between measurement and fit (residuals) as a function of time. (**C**) Histograms of residuals from the low-frequency and high-frequency measurement region (gray and orange, respectively) are compared to our two-region Gaussian error model (black and red).

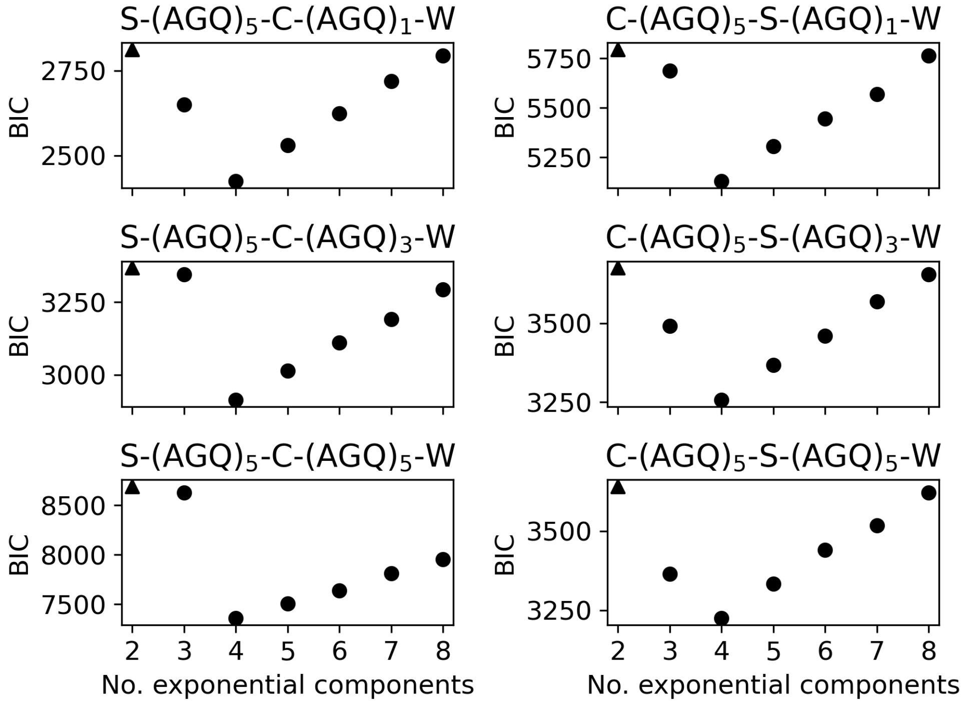


Figure 12: **Identification of optimal number of components for fitting PET measurements.** Bayesian information criterion (BIC) for global fits to experimental data (Fig. 9) with different numbers of exponential components and thereby numbers of parameters, i.e., model complexity. Lower BIC implies a more favorable trade-off between goodness of fit and model complexity.

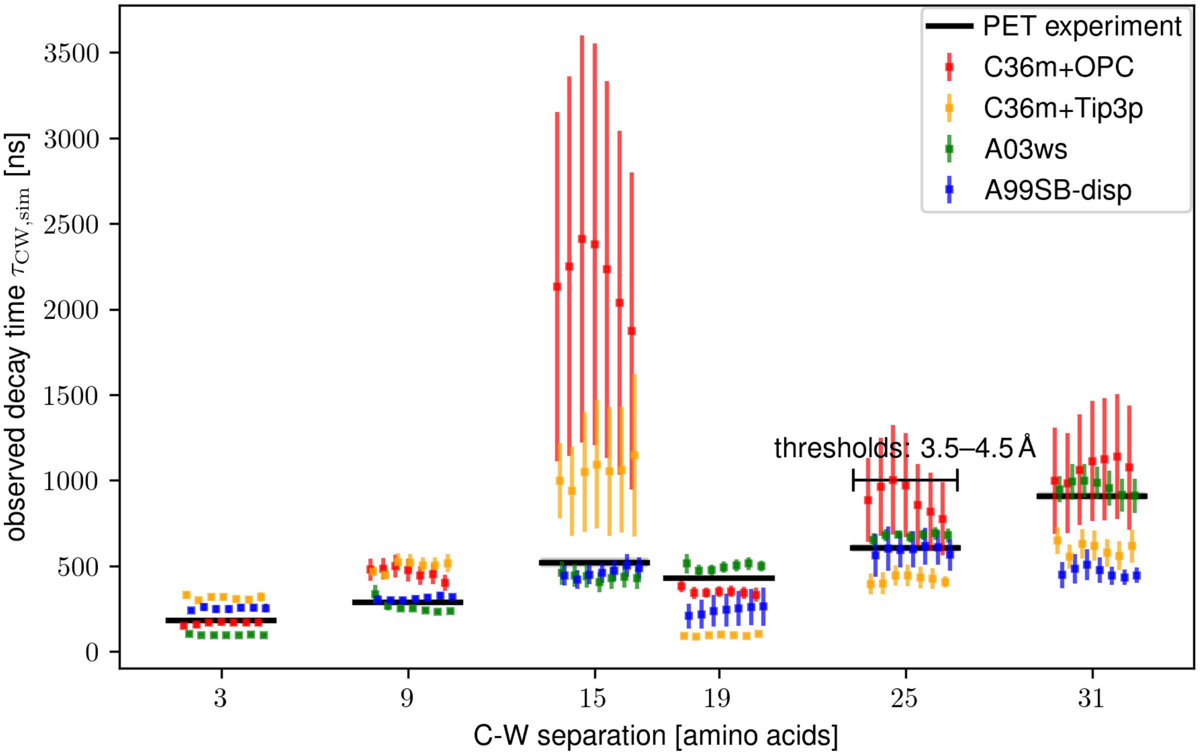


Figure 13: **Effect of contact threshold distance on decay time.** Predictions of observed C-W contact decay times of AGQ-repeat peptides for different C-W contact tresholds $\left(3.5, 3.75, .3.9, 4, 4.1, 4.25, 4.5\,\text{Å}\right)$. All tested C-W separations are indicated by numbers on the x-axis; additional slight horizontal displacement indicates different contact thresholds, increasing from left to right. Errors were estimated by bootstrapping trajectories and represent 68 % confidence intervals.

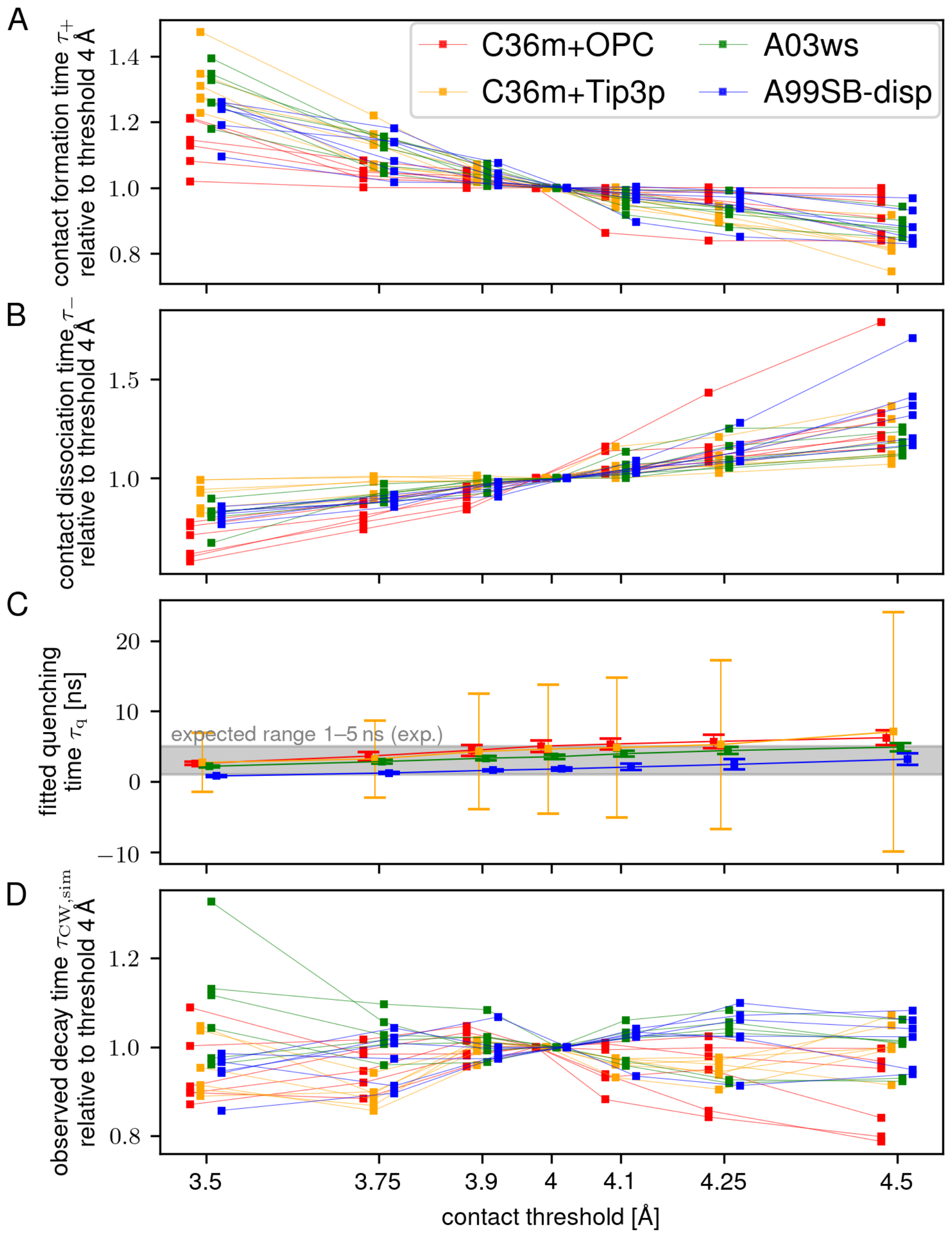


Figure 14: **Transition times of C-W-contact formation kinetics in AGQ-repeat peptides calculated from simulations as a function of contact threshold distance.** Transition times are normalized to their respective estimates for a threshold of 4 Å. All tested contact thresholds are indicated by numbers on the x-axis; additional slight horizontal displacement is for clarity only. The differences in contact formation times ($\tau_+$, panel **A**) and contact dissociation times ($\tau_-$, panel **B**) are mainly compensated for by a different fitted quenching times ($\tau_q$, panel **C**), leading to similar predictions for the observed decay time ($\tau_{CW,sim}$, panel **D**). The fitted quenching times for all tested contact thresholds of 3.5-4.5 Å all fall within the sensible range of 1 ns < $\tau_q$ < 5 ns estimated from experiments (gray area in panel C). Errors in panel C were estimated by bootstrapping trajectories and represent 68 % confidence intervals.

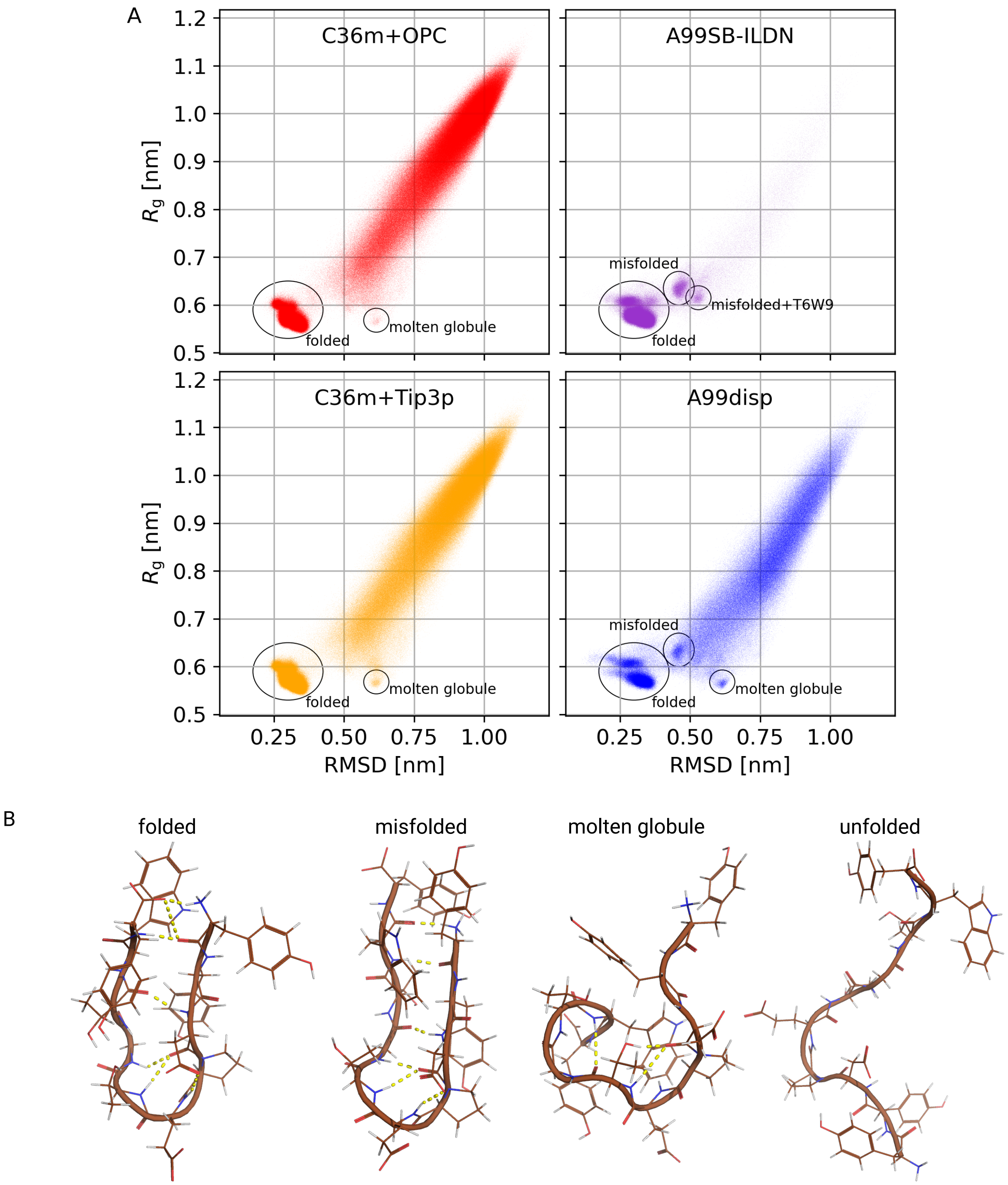


Figure 15: **Comparison of CLN025 ensembles obtained from different force fields.** (**A**) Projection of CLN025 MD trajectories onto two observables, RMSD to the crystal structure and radius of gyration, with conformational states indicated by black ellipses. Unmarked states are unfolded. (**B**) Selected reference structures of CLN025 for the four main conformations found in simulations using A99disp.

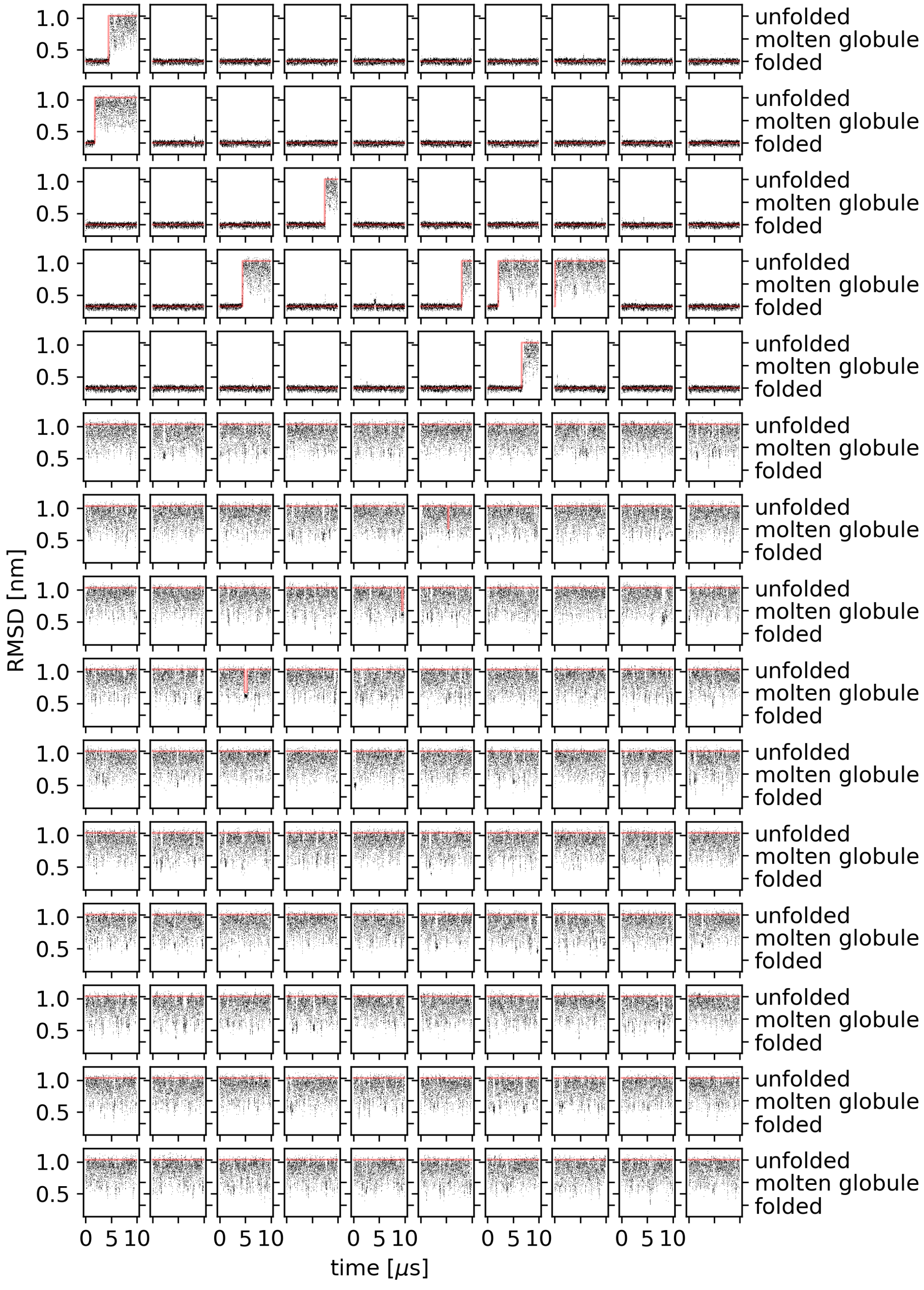


Figure 16: **Assignment of unfolded, molten globular, and folded states of CLN025 in trajectories obtained using C36m+OPC.** Time series of the RMSD to the crystal structure obtained from 150 independent simulations. The red line indicates the assigned conformational state.

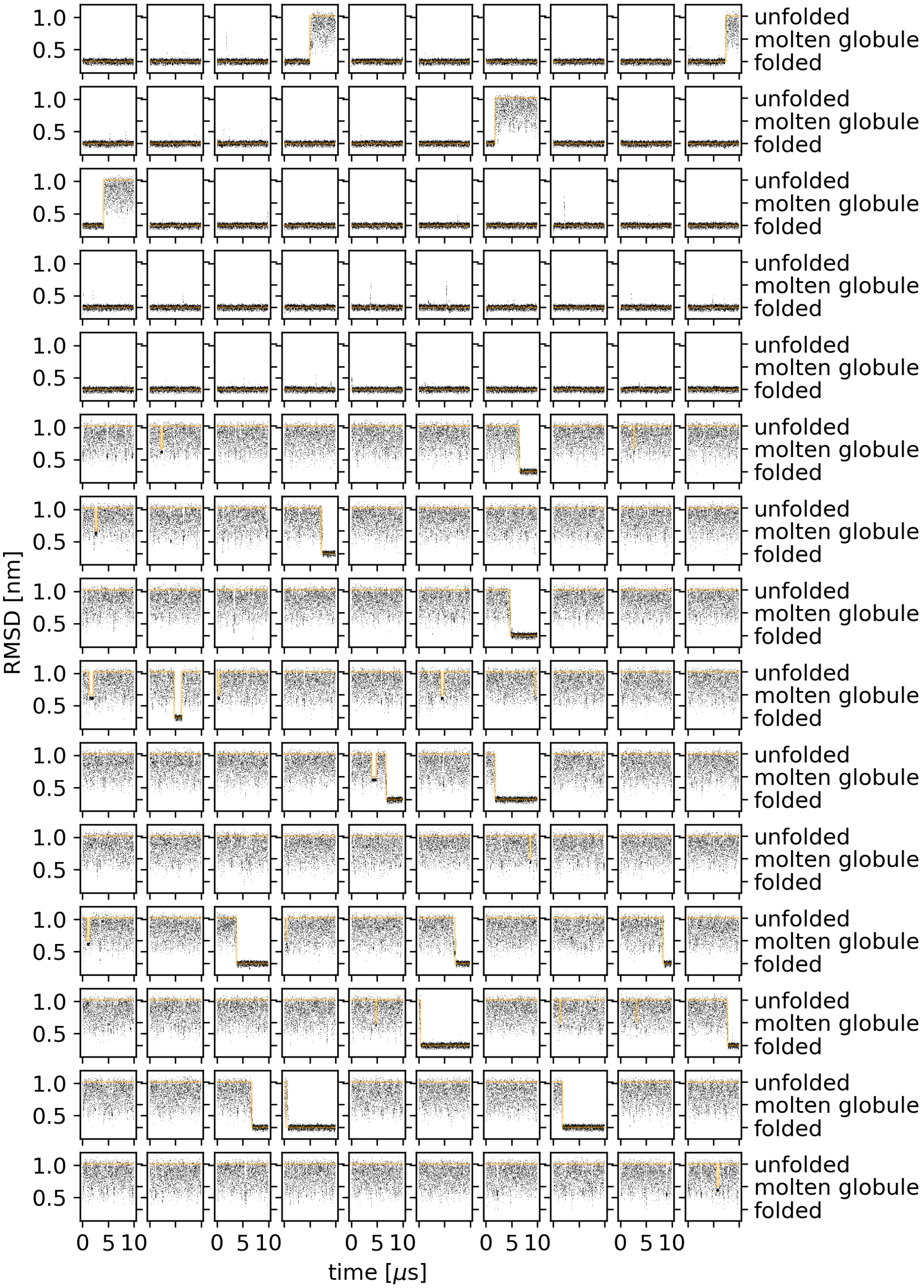


Figure 17: **Assignment of unfolded, molten globular, and folded states of CLN025 in trajectories obtained using C36m+Tip3p.** Time series of the RMSD to the crystal structure obtained from 150 independent simulations. The orange line indicates the assigned conformational state.

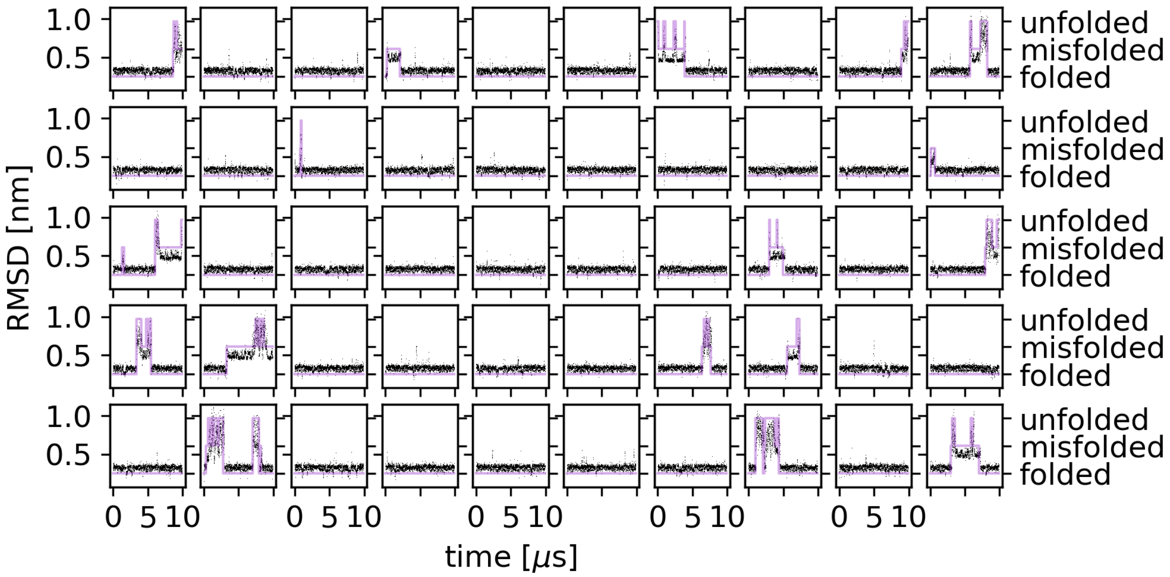


Figure 18: **Assignment of unfolded, misfolded, and folded states of CLN025 in trajectories obtained using A99ildn+Tip4p-Ew.** Time series of the RMSD to the crystal structure obtained from 50 independent simulations. The purple line indicates the assigned conformational state.

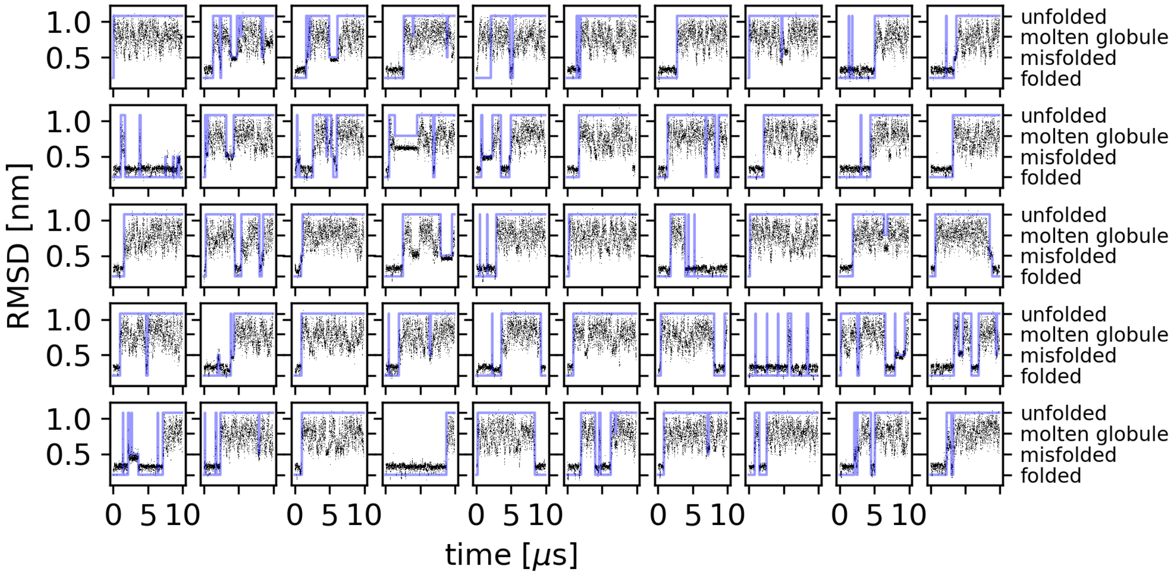


Figure 19: **Assignment of unfolded, molten globular, misfolded, and folded states of CLN025 in trajectories obtained using A99disp.** Time series of the RMSD to the crystal structure obtained from 50 independent simulations. The blue line indicates the assigned conformational state.

Table 1: MD simulation parameters

| Parameter | Value |
|---|---|
| GROMACS version, IDPs & AGQ / CLN025 & glob. | see table 3 / 2018.7 |
| Energy minimization algorithm | steepest descent |
| Energy minimization steps | $\leq 5 \times 10^4$ |
| Equi. length NVT, IDPs & AGQ / CLN025 & glob. | 0.1 ns / 0.5 ns |
| Equi. length NPT 1, IDPs / AGQ | 0.1 ns / 0.1 ns |
| Equi. length NPT 2, IDPs / AGQ | 0.5 ns / 0.5 ns |
| Equi. length NPT, CLN025 / glob. | 1 ns / 1 ns |
| Production run length | see Tables 3 and 2 |
| Position restraints | EM, equi. NVT, equi. NPT 1 |
| Position restraints froce constant | $1000\,\mathrm{kJ\,mol^{-1}\,nm^{-2}}$ |
| Time step (CLN025 and glob.), equi. / prod. | 2 fs / 4 fs |
| Time step (IDPs and AGQ), equi. / prod. | 4 fs / 4 fs |
| Output step, (CLN025, XPCB, and nNOS) | 10 ps |
| Output step, (RNaseA, HEWL, and TIM) | 100 ps |
| Output step, (IDPs and AGQ) | see Table 3 |
| Box shape, IDPs and AGQ / CLN025 and glob. | dodecahedral / triclinic |
| Box extension beyond protein, extended / glob. | > 2 nm / 1.5 nm |
| Salt concentration | $0.15\,\mathrm{mol\,L^{-1}}$ |
| Thermostat | v-rescale[1] |
| Temperature $T$ (CLN025 and glob.) | 300 K |
| Temp. $T$ (IDPs and AGQ), CHARMM / Amber | 300 K / 298 K |
| Temperature coupling constant $\tau_T$ | 0.1 ps |
| Barostat, equi. / prod. and equi. 2 for IDPs and AGQ | Bernedsen[2] / Parrinello-Rahman[3] |
| Pressure $p$ | 1 bar |
| Pressure coupling constant $\tau_p$ (CLN025 and glob.) | 1 ps |
| $\tau_p$ (IDPs and AGQ), C22*/ other CHARMM / Amber | 2 ps / 30 ps / 20 ps |
| Constraints | all bonds |
| Constraint algorithm, solvent / solute | SETTLE[4] / LINCS[5] |
| LINCS order, EM / equi. / prod. | 4 / 4 / 6 |
| Virtual sites[6] | hydrogen atoms |
| Coulomb force algorithm | Particle Mesh Ewald[7] |
| PME order | 4 |
| PME short range cutoff (CLN025 and glob.) | 1 nm |
| PME cutoff (IDPs and AGQ), CHARMM / Amber | 0.95 nm / 1 nm |
| van der Waals force algorithm | cutoff |
| van der Waals cutoff (CLN025 and glob.) | 1 nm |
| vdW cutoff (IDPs and AGQ), CHARMM / Amber | 0.95 nm / 1 nm |
| Dispersion correction | energy and pressure |

Table 2: MD simulation lengths for globular proteins and CLN025

| protein | C36m+OPC | C36m+TIP3P | A99disp | A99SB-ILDN |
|---|---|---|---|---|
| TIM | $20 \times 5$ µs | $20 \times 5$ µs | $20 \times 5$ µs | $20 \times 5$ µs |
| HEWL | $20 \times 5$ µs | $20 \times 5$ µs | $20 \times 5$ µs | $20 \times 5$ µs |
| RNaseA | $20 \times 5$ µs | $20 \times 5$ µs | $20 \times 5$ µs | $20 \times 5$ µs |
| nNOS | $50 \times 1$ µs | $50 \times 1$ µs | $50 \times 1$ µs | $50 \times 1$ µs |
| XPCBD | $50 \times 1$ µs | $50 \times 1$ µs | $50 \times 1$ µs | $50 \times 1$ µs |
| CLN025 | $50 \times 10$ µs $+100 \times 10$ µs $^{(a)}$ | $50 \times 10$ µs $+100 \times 10$ µs $^{(a)}$ | $50 \times 10$ µs | $50 \times 10$ µs |

$^{(a)}$ started from different unfolded structures.

Table 3: MD simulations for IDPs and AGQ peptides

| force field | MeV-Ntail | p53-TAD | CBP-NCBD | p53-pTAD | RSP-8 | AGQ-pept. |
|---|---|---|---|---|---|---|
| | | | Simulation length | | | (6 diff. pept.) |
| C36m+OPC | (6 rep.) 43 µs | 20 × 10 µs | 20 × 5 µs | 5 × 4.7 µs | 5 × 5 µs | 6 × 5 × 5 µs |
| C36m+TIP3P | 5 × 2.5 µs | 20 × 10 µs | 20 × 5 µs | 5 × 2.1 µs | 5 × 5 µs | 6 × 5 × 5 µs |
| C36m+TIP4P | 3 × 5 µs | | | | 1 × 10 µs | |
| C22* | (3 rep.) 17 µs | 20 × 20 µs | (5 rep.) 44.9 µs | 4 × 7 µs | 1 × 10 µs | |
| A03ws | 3 × 5 µs | 24 × 1 µs | 4 × 5 µs | 1 × 10 µs | 1 × 10 µs | 6 × 5 × 5 µs |
| A99disp | 1 × 30 µs | 24 × 4.5 µs | 20 × 5 µs | 5 × 6 µs | 5 × 10 µs | 6 × 5 × 5 µs |
| A99SB-ws | 5 × 4.5 µs | 30 × 20 µs | (20 rep.) 103 µs | 5 × 5 µs | 5 × 5 µs | |
| | | | Output time step | | | |
| C36m+OPC | 1 ns | 10 ns | 10 ns | 1 ns | 10 ns | 1 ns |
| C36m+TIP3P | 5 ns | 10 ns | 10 ns | 1 ns | 10 ns | 1 ns |
| C36m+TIP4P | 8 ns | | | | 20 ns | |
| C22* | 1 ns | 10 ns | 4 ns | 4 ns | 10 ns | |
| A03ws | 2 ns | 5ns | 4 ns | 4 ns | 5 ns | 1 ns |
| A99disp | 2.5 ns | 10 ns | 5 ns | 10 ns | 4 ns | 1 ns |
| A99SB-ws | 1 ns | 50 ns | 5 ns | 5 ns | 5 ns | |
| | | | GROMACS version | | | |
| C36m+OPC | 2019.6 | 2019.6 | 2019.6 | 2019.6 | 2019.6 | 2019.6 |
| C36m+TIP3P | 5.0.8 | 2019.3 | 2019.6 | 2019.6 | 5.0.8 | 2019.6 |
| C36m+TIP4P | 5.0.8 | | | | 5.0.8 | |
| C22* | 2019.6 | 2019.3 | 5.0.8 | 5.0.8 | 4.5.4 | |
| A03ws | 2019.6 | 2019.6 | 5.0.8 | 5.0.8 | 5.0.8 | 2019.6 |
| A99disp | DESMOND[8] | 2019.6 | 2019.6 | 2019.6 | 2019.6 | 2019.6 |
| A99SB-ws | 2019.6 | 2019.3 | 2019.6 | 2019.6 | 2019.6 | |

Format for simulation lengths: number of replica × replica length.
For sets of replica with different lengths: (number of replica) total length.

Table 4: Best-fit parameters for measured PET relaxation curves for all peptides. Amplitudes are reported as mean across measured curves.

| Peptide | $\tau_{\mathrm{in}}$ [ps] | $\tau_1 = \tau_{\mathrm{CW,exp}}$ [ns] | $\tau_2$ [$\mu$s] | $\tau_3$ [$\mu$s] | $\tau_4$ [ms] |
|---|---|---|---|---|---|
| S-(AGQ)$_5$-C-(AGQ)$_1$-W | 198 | 179 | 16.4 | 144 | 10.1 |
| S-(AGQ)$_5$-C-(AGQ)$_3$-W | 216 | 285 | 10.2 | 233 | 9.5 |
| S-(AGQ)$_5$-C-(AGQ)$_5$-W | 210 | 519 | 9.8 | 284 | 9.9 |
| C-(AGQ)$_5$-S-(AGQ)$_1$-W | 230 | 428 | 10.8 | 301 | 9.7 |
| C-(AGQ)$_5$-S-(AGQ)$_3$-W | 186 | 604 | 9.3 | 348 | 10.2 |
| C-(AGQ)$_5$-S-(AGQ)$_5$-W | 216 | 908 | 11.3 | 619 | 16.2 |
| Peptide | $t_{\mathrm{on}}$ [ns] | $A_1$ | $A_2$ | $A_3$ | $A_4$ |
| S-(AGQ)$_5$-C-(AGQ)$_1$-W | 117 | 11812 | 3515 | 2287 | 1649 |
| S-(AGQ)$_5$-C-(AGQ)$_3$-W | 117 | 11250 | 4596 | 2377 | 1606 |
| S-(AGQ)$_5$-C-(AGQ)$_5$-W | 117 | 8735 | 6857 | 2466 | 1612 |
| C-(AGQ)$_5$-S-(AGQ)$_1$-W | 117 | 13155 | 2803 | 1687 | 1529 |
| C-(AGQ)$_5$-S-(AGQ)$_3$-W | 117 | 14388 | 3793 | 2072 | 2008 |
| C-(AGQ)$_5$-S-(AGQ)$_5$-W | 117 | 11181 | 4972 | 1523 | 1829 |

Table 5: Contact formation and dissociation times from simulations.

| Peptide | | C36m+OPC | C36m+TIP3P | A03ws | A99disp |
|---|---|---|---|---|---|
| S-$(AGQ)_5$-C-$(AGQ)_1$-W | $\tau_+$ [ns] | 52 ± 7 | 69 ± 5 | 30.0 ± 2.3 | 131 ± 13 |
| | $\tau_-$ [ns] | 2.22 ± 0.10 | 1.267 ± 0.031 | 1.69 ± 0.07 | 2.01 ± 0.17 |
| S-$(AGQ)_5$-C-$(AGQ)_3$-W | $\tau_+$ [ns] | 194 ± 27 | 109 ± 11 | 110 ± 6 | 158 ± 11 |
| | $\tau_-$ [ns] | 3.5 ± 0.8 | 1.238 ± 0.028 | 2.7 ± 1.0 | 1.91 ± 0.28 |
| S-$(AGQ)_5$-C-$(AGQ)_5$-W | $\tau_+$ [ns] | 750 ± 370 | 230 ± 80 | 151 ± 22 | 214 ± 27 |
| | $\tau_-$ [ns] | 2.32 ± 0.28 | 1.25 ± 0.06 | 2.10 ± 0.20 | 1.58 ± 0.06 |
| C-$(AGQ)_5$-S-$(AGQ)_1$-W | $\tau_+$ [ns] | 122 ± 12 | 26.4 ± 1.8 | 160 ± 11 | 130 ± 60 |
| | $\tau_-$ [ns] | 2.72 ± 0.29 | 1.73 ± 0.11 | 1.69 ± 0.15 | 2.11 ± 0.18 |
| C-$(AGQ)_5$-S-$(AGQ)_3$-W | $\tau_+$ [ns] | 280 ± 90 | 115 ± 16 | 188 ± 10 | 300 ± 60 |
| | $\tau_-$ [ns] | 2.07 ± 0.20 | 1.61 ± 0.07 | 1.38 ± 0.06 | 1.80 ± 0.18 |
| C-$(AGQ)_5$-S-$(AGQ)_5$-W | $\tau_+$ [ns] | 280 ± 100 | 186 ± 27 | 289 ± 28 | 260 ± 50 |
| | $\tau_-$ [ns] | 1.74 ± 0.11 | 2.00 ± 0.09 | 1.47 ± 0.13 | 2.19 ± 0.29 |

### Supplementary Methods

#### p53-pTAD (and other IDPs) CD experiments

Circular Dichroism (CD) spectra were recorded on the AU-CD beamline of the ASTRID2 synchrotron radiation source, Storage Ring facilities (ISA), Aarhus University, Denmark. The spectra were measured at 25 °C using 0.1 mm quartz cuvette under nitrogen atmosphere. The lyophilized recombinant protein sample was dissolved in 10 mM sodium-phosphate buffer (pH = 7.2) with 50 mM NaF. The protein concentration (70 µM) was calculated from sample UV absorption at 280 nm. The final CD spectrum was calculated as the smoothed average of five independently measured and baseline corrected spectra recorded between 178-269 nm. The CD intensities were recorded at every 1 nm, with an average of 15 recordings per measurement per wavelength.

#### p53-pTAD (and other IDPs) SAXS experiments

Small-angle X-ray scattering data were recorded at the European Synchrotron Radiation Facility (ESRF), Grenoble, France, at the BioSAXS beamline BM29 in 2018. Measurements were carried out under sample flow to reduce the radiation damage artifacts. SAXS data were collected over ten measurement windows of 0.3 s. The measured scattering curves were normalized to account for protein concentration, corrected for the buffer signal, and averaged over the ten windows to obtain a final scattering curve. Data were processed using the Edna software package.[9] Measurement conditions were similar to the CD experiment, but protein concentration was adjusted to 280 µM. To monitor aggregation effects, replicate measurements were carried out at protein concentrations 562 µM and 1125 µM. Measurements of p53-pTAD at 1125 µM indicated slight aggregation (likely due to radiation damage).

#### Starting structures for IDP and AGQ-peptide simulations

##### RSP-8

The simulation trajectory of RSP-8 using C22* was provided by S. Rauscher. The A03ws simulation was started from an extended conformation. Simulations trajectories using C36m+OPC, C36m+TIP3P, A99disp, and A99SB-ws were started from 5 peptide conformations. These included one fully extended conformation, and four taken from the C22*trajectory at intervals of 2 µs.

##### p53-pTAD

Two p53-pTAD simulations using C22* were started from structures taken from a p53-TAD/CBP-NCBD complex (PBD ID: 2L14,[10] first an second structure from the NMR ensemble (2L14/1 and 2L14/2)). Two additional simulations were started from extended p53-pTAD structures generated using PyMOL (v2.7).[11] The simulation using A03ws was started from a structure taken from a complex with CBP-NCBD (PDB ID: 2L14/1). For C36m+OPC, C36m+TIP3P, A99disp, and A99SB-ws, simulations were started from five different conformations: a compact one (PDB ID: 2L14/1), two extended structures (the ones also used with C22*), and two partially unfolded structures taken from the C22* trajectory started from the compact structure (PDB ID: 2L14/1) at times 3.5 µs and 5.0 µs.

##### p53-TAD

Trajectories of p35-TAD obtained with C22* and A99SB-ws were kindly provided by R. Klement. To generate twenty partially collapsed starting structures, p35-TAD was simulated in implicit solvent (see below), starting from an extended structure. Simulations using C36m+OPC and C36m+TIP3P were started from the same twenty starting structures. Simulations using A03ws and A99disp also used the same 20 starting structures, but additionally used two extended and one more partially extended starting structures from implicit solvent simulations, and one collapsed conformation taken from preliminary simulations of the p53-TAD/CBP-NCBD complex based on the resolved NMR structure with P53-pTAD (PDB ID: 2L14/1).

##### Implicit solvent simulations of p53-TAD

Implicit solvent MD simulations were carried out for p53-TAD starting from a fully extended conformation, using GROMACS 5.0.8[12] simulation package, the CHARMM force field and a generalized Born (GB) and solvent accessible surface area implicit solvation model.[13] Electrostatic solvation free energies were calculated by a Still Algorithm,[14] a solvent dielectric constant of 80, and an analytical continuum electrostatics calculation scheme for computing solvent accessible surface area for the protein. Neighbor search cut-offs for computing electrostatic, van der Waals, and GB interactions were uniformly set to 1.4 nm. Simulations were carried out for 100 ps to increase the compactness of the protein. The disordered structure of p53-TAD collapsed into a globule in approximately 65 ps, and conformations after 25 ps, 30 ps, and 50 ps were used as starting structures for all-atom explicit solvent MD simulations.

##### CBP-NCBD

Simulations of CBP-NCBD using C22*were started from five different conformations. Three of these were compact structures with high alpha-helical content taken from PDB entries of complexes of CBD-NCBD and different proteins: 2L14[10] (in complex with p53-pTAD), 1KBH[15] (in complex with transcription coactivator ACTR-AD1), and 1ZOQ[16] (in complex with interferon regulatory factor 3). The remaining two conformations were generated by 1 ns implicit solvent simulations started from an extended unfolded structure (with identical parameters as described for p53-TAD). The structure of CBP-NCBD did not completely collapse in this simulation, and structures were taken at 0.5 ns and 1 ns. Simulations using A03ws were started from one compact (PDB ID: 1KBH[15]) and one extended structure (extracted from an implicit solvent simulation at 1 ns). Simulations using C36m+OPC,

C36m+TIP3P, A99disp, and A99SB-ws were started from 20 conformations available at the protein ensemble database[17] (PED, PED00228[18]). These conformations were based on previous ensembles from multiple MD simulations that were refined against CD, SAXS, and NMR measurements.[18]

##### MeV-Ntail

Eight MeV-Ntail conformations were used to start simulations using the various force fields in this work. Starting structures 1-4 were taken from the PED (PED:00020,[18] structures 1, 201, 301, and 601). These conformations only included protein backbone atoms and side chains were added using the autopsf function of VMD[19] (v1.9.3) to obtain all-atom conformations. The fifth structure was produced by an implicit-solvent simulation started from a fully extended all-atom structure created using PyMol[11] (v2.7). The setup of this simulation was identical to implicit-solvent simulations of p53-TAD, and a partially collapsed structure after 47 ps was taken as the fifth starting structure. Starting structures 6–8 were obtained by performing explicit-solvent MD simulations on structures 2–4 using C36m+TIP3P. Representative structures were selected after 646 ns, 1179 ns, and 2095 ns for simulations starting from structures 2, 3, and 4, respectively. Simulations using C22* were started from structures 1, 3, and 5. Simulations using C36m+TIP3P and A99SB-ws were started from structures 1-5. Simulation using C36m+OPC were started from structures 1, 3, and 5-8. Simulations using A03ws were started from structures 3, 6, and 7. The trajectory using A99disp was kindly provided by S. Piana[20] and was started from an extended conformation.

##### AGQ peptides

Starting structures for the six AGQ repeat peptides were generated using seeding simulations. First, initial, fully extended structures of C-$(AGQ)_5$-S-$(AGQ)_{1,3,5}$-W were generated. The structure of C-$(AGQ)_5$-S-$(AGQ)_5$-W was partially collapsed by a stochastic dynamics simulation performed in vacuum (described below). Then, the structures were solvated (150 mM

NaCl), equilibrated, and 6 µs all-atom MD simulations were performed using C36m+OPC, A99disp, and A03ws. To select a representative set of starting structures, $k$-means clustering was performed on C$\alpha$ coordinates with $k = 5$, and the five cluster centers were used as representative starting structures. The five selected starting structures for C-(AGQ)$_5$-S-(AGQ)$_{1,3,5}$-W peptides were mutated to swap the serine and cysteine positions to generate starting structures for the respective other peptides.

##### Stochastic dynamics simulations

A stochastic dynamics simulation was performed on the C-(AGQ)$_5$-S-(AGQ)$_5$-W peptide to reduce its initial extension. It used GROMACS 2019, the C36m force field for protein-protein interactions, inverse friction coefficient $\tau_t = 0.1$ ps, and a reference temperature of 300 K. Van der Waals and Coulomb interactions were calculated using a group-based pair-list scheme with a 1.4 nm cutoff. The simulation was 1 ns long with a 1 fs time step. The end structure was used as a starting structure for explicit-solvent MD simulations.
